# Ultra-low-field brain MRI morphometry: test-retest reliability and correspondence to high-field MRI

**DOI:** 10.1101/2024.08.14.607942

**Authors:** František Váša, Carly Bennallick, Niall J. Bourke, Francesco Padormo, Levente Baljer, Ula Briski, Paul Cawley, Tomoki Arichi, Tobias C. Wood, David J. Lythgoe, Flavio Dell’Acqua, Thomas C. Booth, Ashwin V. Venkataraman, Emil Ljungberg, Sean C.L. Deoni, Rosalyn J. Moran, Robert Leech, Steven C.R. Williams

## Abstract

Magnetic resonance imaging (MRI) enables non-invasive monitoring of healthy brain development and disease. Widely used higher field (*>*1.5 T) MRI systems are associated with high energy and infrastructure requirements, and high costs. Recent ultra-low-field (*<*0.1T) systems provide a more accessible and cost-effective alternative. However, it remains uncertain whether anatomical ultra-low-field neuroimaging can be used to reliably extract quantitative measures of brain morphometry, and to what extent such measures correspond to high-field MRI. Here we scanned 23 healthy adults aged 20-69 years on two identical 64 mT systems and a 3 T system, using T_1_w and T_2_w scans across a range of (64 mT) resolutions. We segmented brain images into 4 global tissue types and 98 local structures, and systematically evaluated between-scanner reliability of 64 mT morphometry and correspondence to 3 T measurements, using correlations of tissue volume and Dice spatial overlap of segmentations. We report high 64 mT reliability and correspondence to 3 T across 64 mT scan contrasts and resolutions, with highest performance shown by combining three T_2_w scans with low through-plane resolution into a single higher-resolution scan using multi-resolution registration. Larger structures show higher 64 mT reliability and correspondence to 3 T. Finally, we showcase the potential of ultra-low-field MRI for mapping neuroanatomical changes across the lifespan, and monitoring brain structures relevant to neurological disorders. Raw images are publicly available, enabling systematic validation of pre-processing and analysis approaches for ultra-low-field neuroimaging.

## Introduction

Magnetic resonance imaging (MRI) can provide detailed insight into the anatomy and physiology of a wide range of organs, in health and disease. However, conventional higher field MRI systems (*>*1.5 T) have significant infrastructure requirements (including shielding, stable power and cryogenic cooling) as well as associated installation and running costs.

The recent development of ultra-low-field MRI scanners, such as the 64 mT Hyperfine Swoop, promises to revolutionize neuroimaging (Arnold et al., 2023; Kimberly et al., 2023). Ultra-low-field MRI systems can be run using a standard electrical socket and are portable, enabling scanning at the bedside (Mazurek et al., 2021; Balaji et al., 2024) and in low-and middle-income settings with limited access to MRI (Chetcuti et al., 2022; Abate et al., 2024).

Initial studies have shown good correspondence of quantitative estimates of brain morphometry. Most studies to date relied on synthetic enhancement of ultra-low-field scans using deep-learning image quality transfer. Iglesias et al. (2023) and Islam et al. (2025) reported significant correlations between volumetric measurements from super-resolved adult ultra-low-field and high-field scans, and Sorby-Adams et al. (2024) reported significant volume correlations as well as high Dice overlap of segmentations from super-resolved ultra-low-field and high-field scans of patients with Alzheimer’s disease. Concurrently, both Cooper et al. (2024) and Pretzsch et al. (2025) applied *FreeSurfer* to (distinct) datasets of children and/or young adults, and reported good agreement of tissue volume and surface area, and moderate correspondence of cortical thickness estimates. Finally, Ringshaw et al. (2025) and Hsu et al. (2025) extracted accurate volumes from ultra-low-field scans without deep-learning image enhancement, by combining orthogonal ultra-low-field acquisitions. Other recent studies included concurrent acquisition of paired ultra-low-field and high-field data, across a range of participants including infants (Baljer et al., 2024, 2025) or patients with multiple sclerosis (Arnold et al., 2022); however, these studies were not primarily focused on the correspondence of morphometric estimates. Notably, few of these studies investigated the test-retest reliability of 64 mT imaging and derived morphometric measurements (Sorby-Adams et al., 2024; Hsu et al., 2025), which are crucial to enable pooling of data across sites (Abate et al., 2024). Additionally, no prior studies have publicly shared the underlying MRI data in a readily available form, hindering the development of new approaches for analyzing ultra-low-field MR images.

Here, we present the results of morphometric analyses of healthy adults scanned on two identical 64 mT scanners, and a 3 T scanner, without prior image enhancement using image quality transfer. We scanned 23 healthy adult participants using T_1_w and T_2_w scans, including four different voxel resolutions at ultra-low-field, three of which were combined to generate a fifth higher-resolution volume. We segmented and parcellated all scans into 98 structures across 4 tissue types, and extracted corresponding tissue volumes, using SynthSeg+ (Billot et al., 2023). We used these estimates to quantify between-scanner test-retest reliability of 64 mT scans, and their correspondence to 3 T scans, using both correlations of volume across participants, and Dice overlap of segmentations within participants (following rigid-registration of 64 mT to 3 T scans within participants). For details, see Fig. 1 and Methods. We report excellent between-scanner test-retest reliability of 64 mT scan morphometry, and correspondence to 3 T MRI, across contrasts and resolutions. The underlying raw MRI scans and analysis code are publicly available, to help accelerate the development and optimization of methods for the pre-processing and analysis of ultra-low-field brain MRI data.

**Figure 1:**
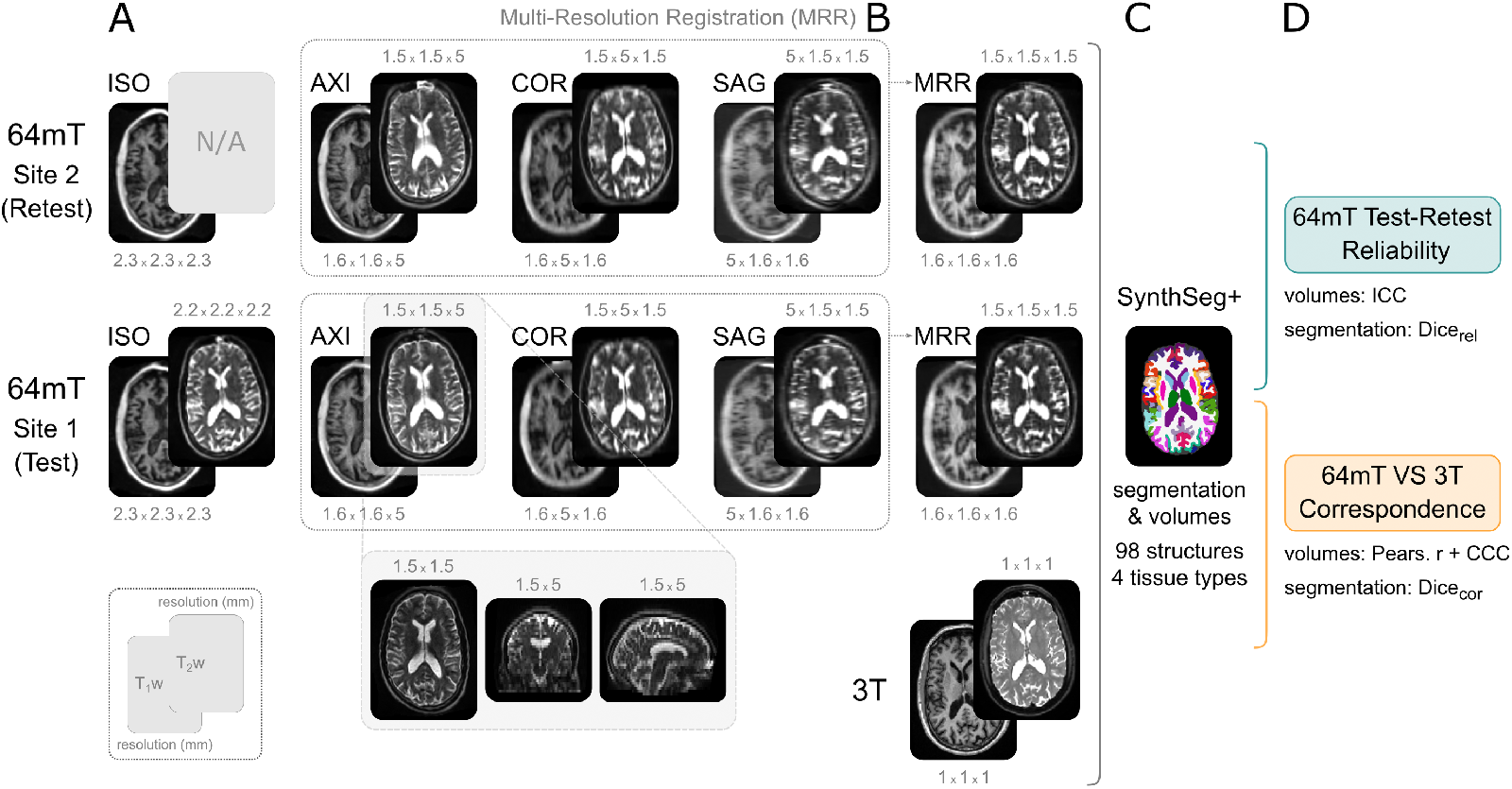
Data and pre-processing, illustrated on one example participant. A) Each participant was scanned on two identical 64 mT scanners and a 3 T scanner, using T_1_w and T_2_w scans, across a range of 64 mT resolutions (listed in mm above or below each scan). The second 64 mT site did not include a T_2_w ISO scan. For details regarding acquisition parameters, see supplementary Table S1. B) The three scans with high in-plane resolutions were combined into a single higher-resolution volume using multi-resolution registration (MRR; Deoni et al., 2022). C) All scans were segmented and parcellated into 98 structures across 4 tissue types using SynthSeg+ (Billot et al., 2023). D) The resulting volumes and segmentations were used to quantify the test-retest reliability of 64 mT scans, and their correspondence to 3 T MRI (ICC: intraclass correlation coefficient; CCC: Lin’s concordance correlation coefficient). Note that most scans depicted here were rigid-registered to the high-field scan of the same contrast, except the bottom panel showcasing all three planes of a “raw”, unregistered, T_2_w axial scan.

## Methods

### Design

In total, 23 healthy adult participants were recruited. To obtain a sample balanced for age and sex, a minimum of 2 male and 2 female participants were recruited from each of the following 5 age-strata: 20-29 years, 30-39 years, 40-49 years, 50-59 years and 60-69 years. Basic demographics including sex, and year and month of birth, were collected. Exclusion criteria were contraindications to MRI (including claustrophobia), major neurological or psychiatric conditions, or other health issues known to affect the brain. Screening was conducted using the standard MRI safety questionnaire and through self-report of medical history. Participants provided informed written consent for all aspects of the study. The study was approved by the King’s College London human Research Ethics Committee (Reference: HR/DP-21/22-26481).

### MRI data acquisition

MRI data for each participant were acquired during two scanning sessions, held no more than 36 days apart. For a distribution of time durations between the two sessions, see supplementary Figure S2. During the first session, at the Centre for Neuroimaging Sciences (CNS; Institute of Psychiatry, Psychology and Neuroscience, King’s College London), participants were scanned for 45 minutes on a GE SIGNA Premier 3 T high-field scanner with a 48-channel head coil (GE Healthcare, Waukesha, WI), followed immediately by a 1-hour protocol on a Hyperfine Swoop 64 mT ultra-low-field scanner (Hyperfine Inc., Guildford, CT). During the second session, at the Evelina Newborn Imaging Centre (ENIC; St. Thomas’s Hospital, Guys and St. Thomas’s NHS Foundation Trust), participants were scanned for 1 hour on a second Hyperfine Swoop system, identical to the first, and running the same software version (rc8.6.0). In all three scanners, Pearltec inflatable pads were used to minimize head motion (Pearl Technology AG, Switzerland).

The 3 T high-field protocol included T_1_w (MPRAGE) and T_2_w scans, acquired at 1*×*1*×*1 mm^3^ resolution. 64 mT protocols included both T_1_w and T_2_w scans, acquired using non-isotropic product sequences (T_1_w: 1.6×1.6×5 mm^3^; T_2_w: 1.5×1.5×5 mm^3^) with high resolution in the axial, sagittal or coronal plane, respectively referred to as AXI, SAG, and COR. Additionally, a research agreement with Hyperfine Inc. enabled acquisition of a custom isotropic sequence (T_1_w: 2.3×2.3×2.3 mm^3^, T_2_w: 2.2×2.2×2.2 mm^3^), referred to as ISO. All scans were acquired on both scanners, except T_1_w ISO scans, which were only acquired during the first scanning session at the CNS (the second scanning session at ENIC instead included an axial T_2_-FLAIR scan of similar duration, not included in these analyses). For further details regarding the sequences, including acquisition times, see supplementary Table S1, main Fig. 1A and supplementary Fig. S8.

### Pre-processing

We used multi-resolution registration (Deoni et al., 2022) to iteratively co-register and average the three orthogonal non-isotropic 64 mT scans (AXI, COR, SAG) into a single “MRR” scan, of a higher effective resolution (T_1_w: 1.6×1.6×1.6 mm^3^, T_2_w: 1.5×1.5×1.5 mm^3^). This was implemented using the *antsMultivariateTemplateConstruction2*.*sh* function from ANTs (Avants et al., 2009) (Fig. 1B). For full details of the MRR pipeline, see the Supplementary Information, and accompanying code. We also note that a similar approach has been described recently Hsu et al. (2025).

We then used SynthSeg+ (Billot et al., 2023), including both “robust” segmentation and cortical parcellation, resulting in 98 regional labels for each scan, and corresponding volume estimates. We note that SynthSeg+ previously demonstrated strong performance on ultra-low-field MRI scans super-resolved using LF-SynthSR (Iglesias et al., 2023; Sorby-Adams et al., 2024; Islam et al., 2025). We additionally grouped most labels into four categories by tissue class, including cortical grey matter (GM_cort_), subcortical grey matter (GM_subc_), white matter (WM) and cerebrospinal fluid (CSF). For details of the names of individual SynthSeg+ structures and their assignment to tissue classes, see supplementary Table S4.

To assess the between-scanner test-retest reliability of 64 mT segmentations and their correspondence to 3 T scans at the voxel level, we rigid-registered each 64 mT scan to the 3 T scan of the corresponding contrast, using FSL FLIRT (Jenkinson and Smith, 2001), and subsequently applied the transform to segmentation outputs (with nearest-neighbour interpolation). This resulted in each participant’s test-retest 64 mT segmentations also being co-registered.

We visually assessed the quality of all individual scans, segmentations and registrations. Raw scans and segmentations were visually of high quality; however, some rigid registrations of 64 mT to 3 T scans appeared imperfect (see supplementary Table S5). In the main text we report results from all scans; for a sensitivity analysis quantifying Dice overlap of segmentations (affected by registration quality) in a subset of scans with high-quality registrations, see supplementary Table S6. We note that approaches for assessing the quality of segmentations and registrations are under constant development and refinement, and there is no consensus reference standard. Combining rigid registration with the Dice coefficient to quantify overlap is an approach frequently used to evaluate performance of neuroimaging segmentation algorithms, both at high-field (e.g., Shattuck et al., 2009; Billot et al., 2023) and ultra-low-field strength (e.g., Sorby-Adams et al., 2024; Hsu et al., 2025). Still, visual assessment of registration quality may be subjective, but will impact Dice scores; as such, Dice-related analyses should be interpreted with caution.

### Analysis

Most analyses were repeated using both 4 “global” aggregate tissue volumes and corresponding segmentation masks, as well as 98 “local” SynthSeg+ structures, on both T_1_w and T_2_w contrasts, across all 64 mT resolutions. Correspondence of 64 mT to 3 T data was assessed using 64 mT scans acquired at the CNS, on the same day as (and immediately after) 3 T scans.

We assessed between-scanner test-retest reliability of volume estimates across participants using the intraclass correlation coefficient ICC(3,1) (Chen et al., 2018), referred to as ICC. We assessed correspondence of volumes from 64 mT and 3 T scans using both Pearson’s r, to quantify linear correlation; and Lin’s concordance correlation coefficient (CCC), to quantify exact agreement (or alignment with the *x* = *y* identity line). We used the Dice overlap of segmented structures within participants to quantify the reliability of 64 mT segmentations (Dice_rel_) and their correspondence to 3 T segmentations (Dice_cor_). We also repeated Dice overlap analyses only within subjects with high-quality rigid registrations of 64 mT to 3 T scans (and consequently also 64 mT scans to each other). The number of participants with high-quality registrations differed by scanner site, contrast and resolution; for numbers of participants across image types, see supplementary Table S5, and for resulting summary statistics, see supplementary Table S6.

To quantify the extent to which 64 mT scans tend to under-or over-estimate tissue volumes relative to 3 T, we quantified the percentage difference in volumes relative to high-field data (supplementary Table S2).

To assess whether larger and more reliable structures show higher correspondence to 3 T scans, we quantified Spearman’s rank correlations between local maps of 64 mT reliability, correspondence to 3 T, and median regional volume from 3 T scans. This analysis was limited to 64 mT T_2_w MRR scans. We also used linear models to estimate changes in tissue volume as a function of age, covarying for sex, using both 3 T and 64 mT scans. We quantified the relationship across field strengths between regional rates of change (t-statistics) and variance explained by age and sex (r^2^) using Pearson’s r and Lin’s CCC (supplementary Table S3).

Finally, we extracted residuals from these linear models, to obtain individual tissue volumes adjusted for effects of participant age and sex (for T_2_w 3 T scans and T_2_w 64 mT MRR scans). We then computed robust-scaled adjusted volumes within each region and across participants, to obtain non-parametric equivalents of the Z-score (Leys et al., 2013), a measure of individual deviation from the norm. We then identified the participant and region showing the greatest deviation from the median in T_2_w high-field scans, and visualized the corrected volume of this region from all three scanners, across participants. Additionally, we quantified the reliability and correspondence of all regional deviations in this participant, using ICC, Pearson’s r and Lin’s CCC.

## Results

### Data acquisition and quality

We recruited 23 healthy adult participants aged 20-69 years, with 2-3 males and 2-3 females per decade (for participant distribution by age and sex, see supplementary Figure S1). We acquired both T_1_w and T_2_w scans on a 3 T high-field scanner (GE Healthcare SIGNA Premier, Waukesha, WI) and two identical 64 mT ultra-low-field scanners (Hyperfine Swoop, Guildford, CT; hardware version: 1.7, software version: 8.6.0).

Ultra-low-field scans were acquired at four resolutions, including three product sequences with higher resolution within the axial, coronal or sagittal plane (1.5×1.5×5 mm^3^ or 1.6×1.6×5 mm^3^, hereafter referred to as AXI, COR, SAG respectively) and a custom isotropic sequence of approximately equivalent voxel volume (2.2 mm^3^ or 2.3 mm^3^ isotropic, hereafter referred to as ISO). We further combined AXI, COR and SAG scans into one within participants using multi-resolution registration (Deoni et al., 2022), yielding a single higher-resolution “MRR” volume per contrast (of effective resolution 1.5 mm^3^ or 1.6 mm^3^ isotropic). High-field scans were acquired at 1 mm^3^ isotropic resolution. For details regarding scan parameters, see supplementary Table S1.

Identical contrasts were acquired on both 64 mT scanners, except for T_2_w ISO scans, which were only acquired at one site and therefore omitted from test-retest analyses. Moreover, the T_1_w SAG scan did not properly reconstruct for one participant, resulting in analyses on T_1_w SAG and T_1_w MRR scans being carried out on a sample of 22 (out of 23) participants.

Ultra-low-field MRI scans are of high quality on visual inspection, particularly within the high-resolution plane of acquisition (supplementary Fig. S8), but also inherently of lower visual quality compared to high-field MRI, characterized by lower resolution, reduced contrast between tissue types and increased noise (Fig. 1A,B). We visually controlled the quality of all raw scans, their SynthSeg+ segmentations (Billot et al., 2023) and FSL FLIRT rigid-registrations within participants (Jenkinson and Smith, 2001). For details, see Methods and Supplementary Information.

Results were broadly consistent across contrasts and 64 mT resolutions. In the main text, we focus primarily on results obtained using T_2_w MRR 64 mT scans, which show the highest between-scanner reliability and correspondence to 3 T scans. We summarize the reliability and correspondence to 3 T for both T_1_w and T_2_w scans and across resolutions in Table 1 and supplementary Figures S4-S7.

**Table 1:**
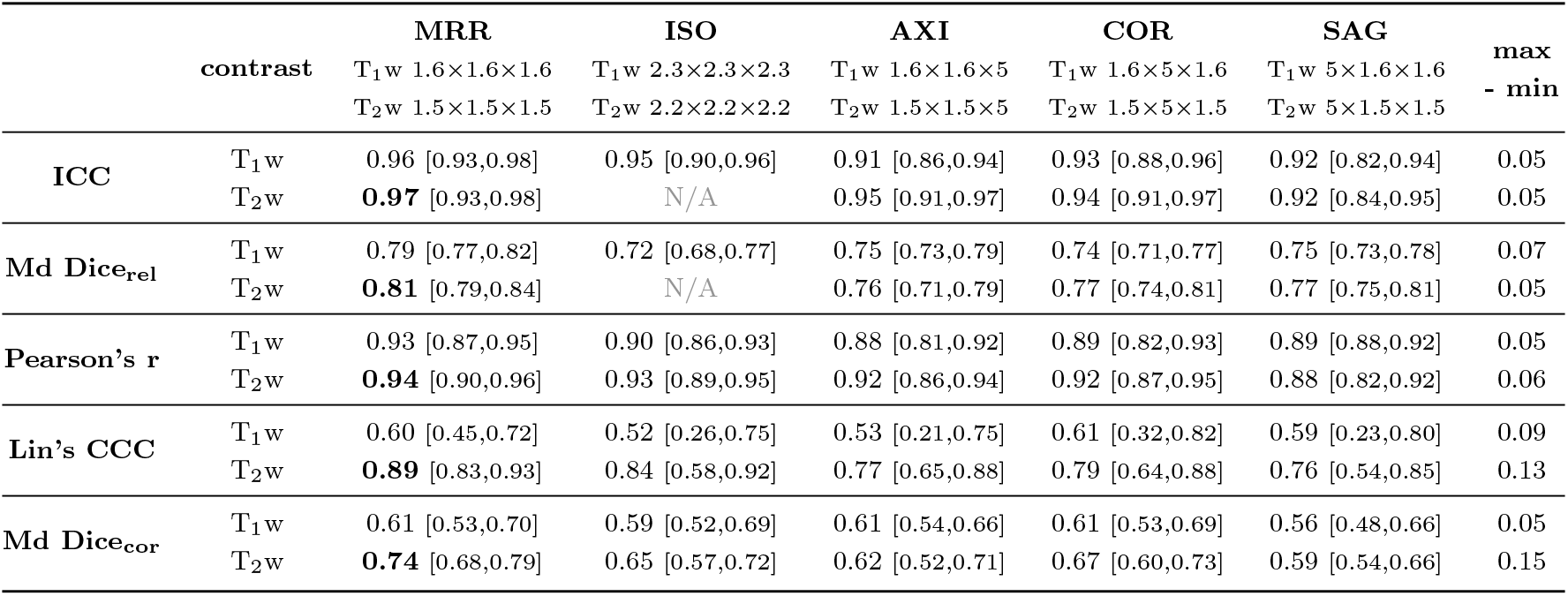
Summary of regional reliability and correspondence, across contrasts and resolutions. Main values correspond to median [first, third quartile], i.e. Md [Q_1_, Q_3_], across 98 SynthSeg+ labels. The last column is the difference between the maximum and minimum Md value within each row. Maximum values for each statistic are highlighted in bold.

### Global reliability of 64 mT and correspondence to 3 T

Ultra-low-field T_2_w MRR scans combined using multi-resolution registration (Deoni et al., 2022) showed excellent reliability between both 64 mT scanners, across all four tissue types (cortical grey matter, GM_cort_; subcortical grey matter, GM_subc_; white matter, WM; cerebrospinal fluid, CSF; for details regarding the assignment of individual structures to tissue types, see supplementary Table S4). Correlations of tissue volumes across participants were particularly high, with intraclass correlation coefficient (ICC) values of 0.98-1.0 (Fig. 2A). Accurate voxel-level labeling of tissues is a more difficult task, with commensurately lower but still very high reliability, including a reliability Dice overlap (referred to as Dice_rel_) of underlying segmentations within participants showing median values of 0.76-0.92 (Fig. 2B).

**Figure 2:**
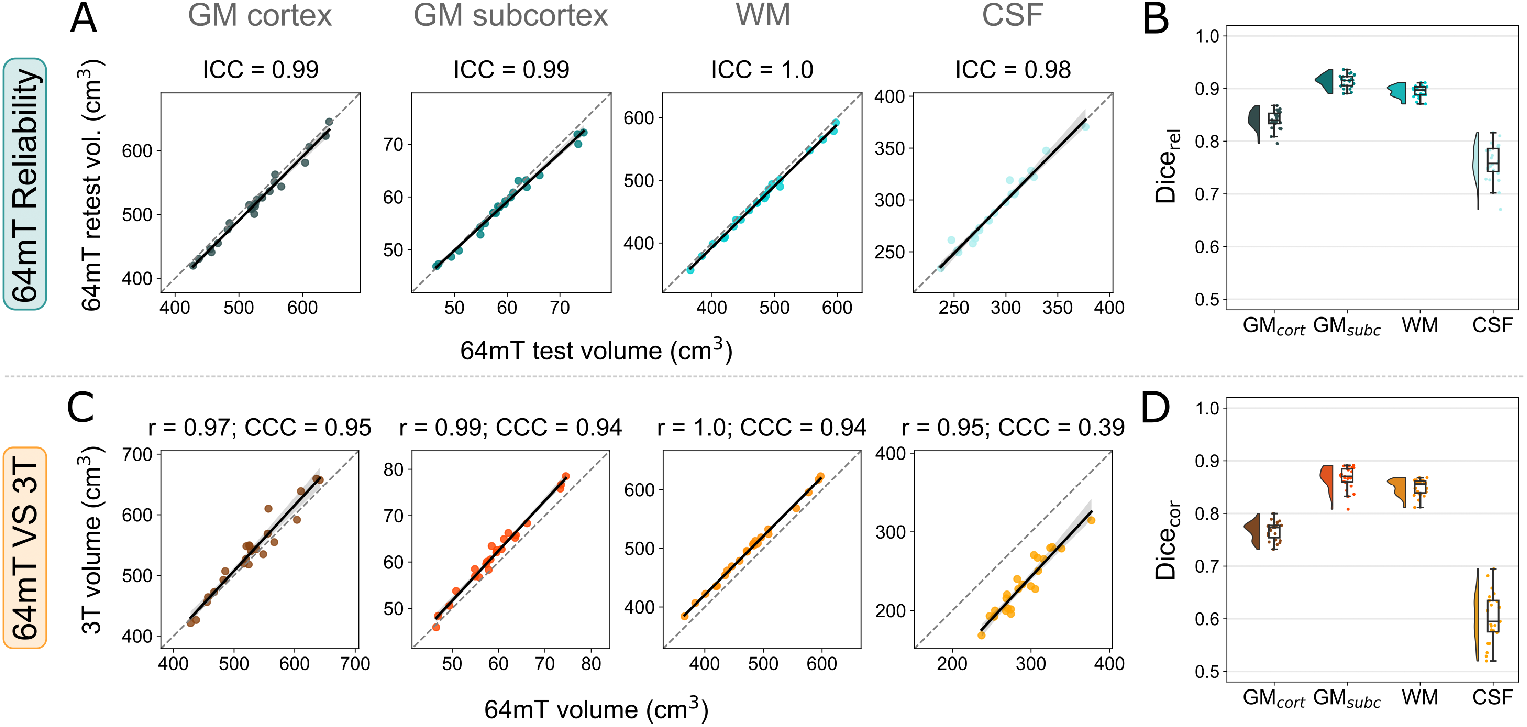
Global reliability of 64 mT scans, and correspondence to 3 T MRI. A) Correlations of global tissue volumes between both 64 mT sites, across participants. B) The Dice overlap of underlying rigid-registered segmentations within participants. C) Correspondence of 64 mT (site 1) data to 3 T, across participants. Statistics indicate the linear correlation (Pearson’s r) and exact agreement, or alignment with the *x* = *y* identity line (Lin’s concordance correlation coefficient; CCC). D) Overlap of segmentations across field strengths, within participants. For results across all contrasts and resolutions, see supplementary Figures S4-S7.

Global tissue volumes extracted from 64 mT T_2_w MRR scans showed both high linear correlations to 3 T counterparts, with Pearson’s r across tissue types in the range 0.95-1.0, but also under-or over-estimates resulting in Lin’s concordance correlation coefficient (CCC), quantifying exact agreement, in the range 0.39-0.95 (Fig. 2C). The correspondence Dice overlap (referred to as Dice_cor_) of globally-aggregated segmentation labels across field strengths was lower than between both 64 mT scanners, but remained high, with median values in the range 0.60-0.87 (Fig. 2D).

For results of global 64 mT reliability and correspondence to 3 T across contrasts and resolutions, see supplementary Figures S4-S7.

Assessment of percentage difference in volume estimates at 64 mT, relative to 3 T, confirmed median underestimates of volume in grey and white matter (GM_cort_ = − 1.5%, GM_subc_ = − 3.7%, WM = − 4.4%), and overestimates in the CSF (+24.6%), apparent in global correspondence plots (Fig. 2C). These differences may be due to partial volume effects, as well as other causes (see Discussion). Individual structures showed a similar trend, with median values ranging from *−*25.7% to +28.0% (Md [Q_1_,Q_3_] = − 1.6 [− 4.5, −0.9]%). For a summary of percentage differences across contrasts and resolutions, see supplementary Table S2.

### Local reliability of 64 mT and correspondence to 3 T

We next repeated analyses locally, across all 98 structures provided by the SynthSeg+ segmentation and cortical parcellation (Billot et al., 2023). In Figures 3 and 4, we show both an overview of reliability and correspondence statistics across regions (left), as well as detailed results for three exemplar regions for each statistic, showcasing structures with minimum, median and maximum values (right).

**Figure 3:**
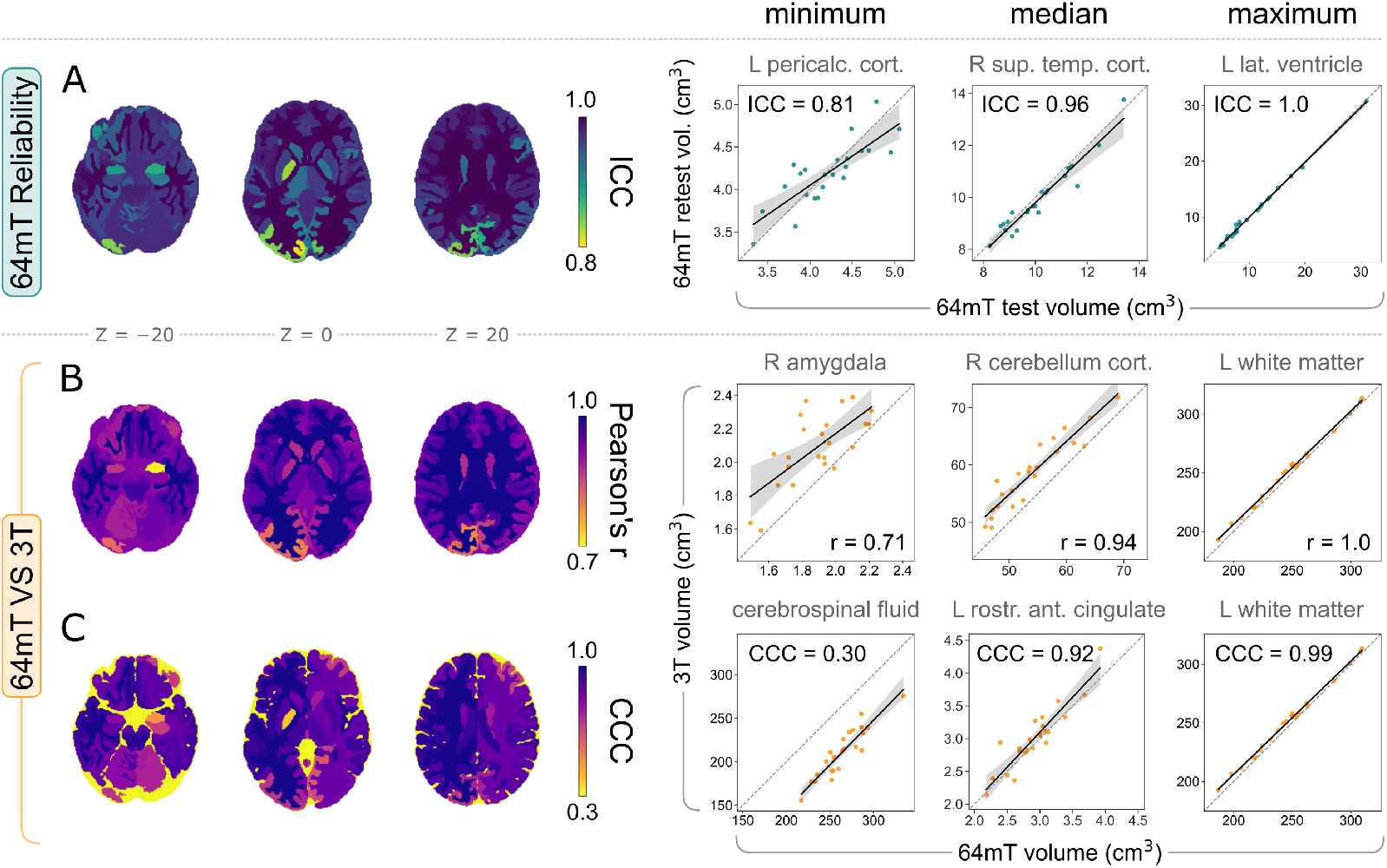
Local reliability of 64 mT tissue volumes, and correspondence to 3 T MRI. Left: summary of correlation statistics across regions, on three axial slices of a segmented high-field MNI template. Right: the corresponding regions with minimum, median and maximum volume correlations. A) 64 mT between-scanner reliability of local volumes, quantified using the intraclass correlation coefficient (ICC). Below, the correspondence of local 64 mT volumes to 3 T, assessed using B) Pearson’s r, which quantifies linear correlation, and C) Lin’s concordance correlation coefficient (CCC), which quantifies exact agreement.

**Figure 4:**
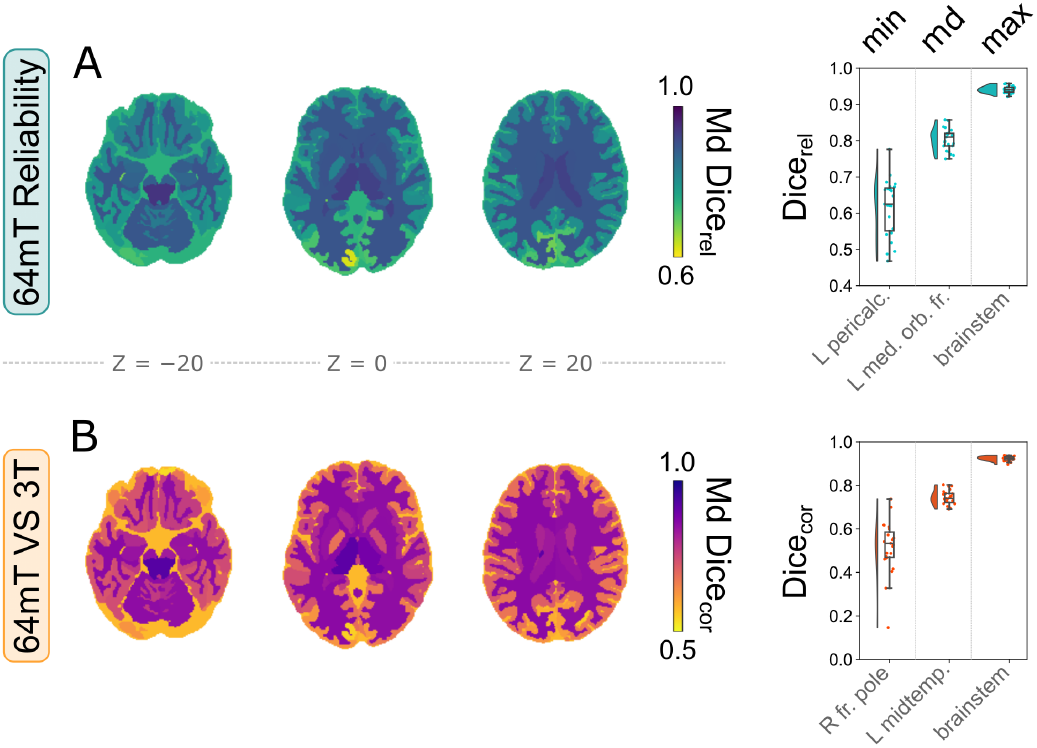
Local reliability of 64 mT segmentations, and correspondence to 3 T MRI. Left: median Dice overlap of all structures across participants, on three axial slices of a segmented high-field MNI template. Right: the corresponding regions with minimum, median and maximum Dice overlap. A) Reliability of local segmentations between 64 mT scanners. B) Correspondence of local segmentations between 64 mT and 3 T scans.

Between-scanner reliability of 64 mT local volumes demonstrated variability across regions, but remained very high overall, with median ICC = 0.96 (Fig. 3A). The correspondence of volumes extracted from 64 mT scans to 3 T counterparts remained high at the individual structure level as well, with both high linear correlations exemplified by median r = 0.94 (Fig. 3B) and good agreement, shown by median CCC = 0.92 (Fig. 3C). However, some structures showed lower values, including for example minimum r = 0.71 and minimum CCC = 0.30, in the right amygdala and extra-cortical CSF respectively (Fig. 3B,C).

Similarly to global tissue types, we next assessed the median (across participants) Dice overlap of segmentations at the local level, both within 64 mT scans (Dice_rel_) and across field strengths (Dice_cor_) (Fig. 4). Segmentations derived from both 64 mT sites showed high overlap, including median Dice_rel_ = 0.81 across regions, but also moderate variability, with a Dice_rel_ coefficient range of 0.63-0.94 (Fig. 4A). Similarly to global volumes, the overlap of 64 mT segmentations to 3 T counterparts was slightly lower (than reliability), but remained high, with median Dice_cor_ = 0.74 and a range of 0.53-0.92 (Fig. 4B), indicating that ultra-low-field scans give rise to generally reliable and accurate voxel-wise segmentations at the local level as well.

### Relationships between regional statistics

We next wished to ascertain whether structures with more reliable 64 mT volume estimates and segmentations show higher correspondence to 3 T scans, and whether these quantities are additionally related to the size of regions, as an extension of prior work showing that larger regions tend to be more reliably estimated in high-field scans (e.g., Morey et al., 2010; Whelan et al., 2016; Madan and Kensinger, 2017; Elliott et al., 2023. Additionally, our key regional statistics (i.e., ICC, Pearson’s r, Lin’s CCC, Dice coefficient) are sensitive to related but different facets of reliability and correspondence, and we wished to quantify the extent of empirical similarities or differences between these metrics. We therefore calculated Spearman’s rank correlations between the 98 regional values of each statistic, across all pairs of six measures, including both measures of 64 mT reliability (ICC and median Dice_rel_), three measures of 3 T correspondence (Pearson’s r, Lin’s CCC and median Dice_cor_) and the median volume of regions across all participants, as estimated using 3 T MRI scans (Fig. 5). Measures were positively correlated, with 7/15 relationships surviving Bonferroni correction for multiple comparisons (i.e. *p <* 0.05*/*15 = 0.0033), indicating that larger structures can be more reliably estimated using 64 mT scans, and show greater correspondence to 3 T scans. Relationships between different measures of 64 mT reliability were strong and relatively linear, as were relationships between measures of 64 mT reliability and correspondence across fields; however, relationships between regional volume and remaining statistics were commensurately weaker and showed nonlinear relationships. For scatterplot illustrations of four exemplar relationships, see Fig. 5B.

**Figure 5:**
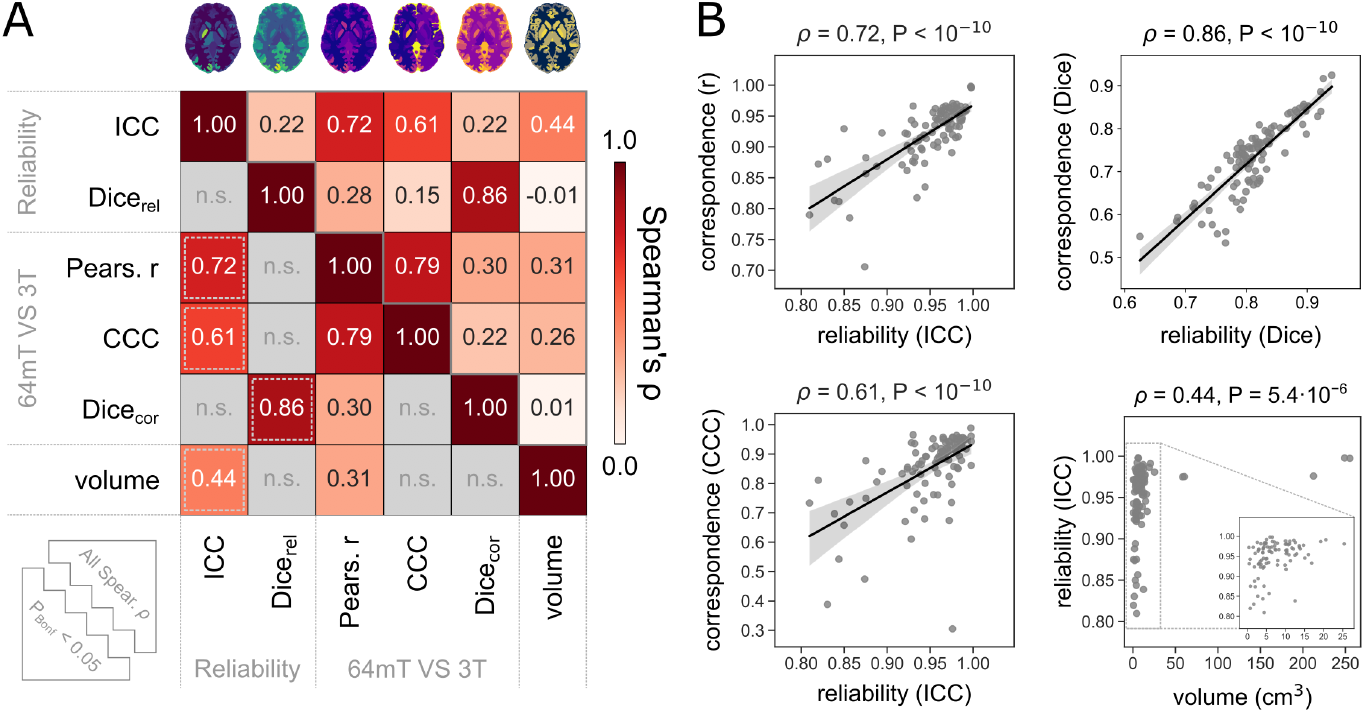
Larger structures are more reliably estimated using 64 mT MRI and show greater correspondence to 3 T MRI. A) Spearman’s rank correlations between local maps, of 64 mT reliability, correspondence to 3 T, and average volume of each structure (extracted from high-field scans). In the lower triangular part, only correlations which remain significant after Bonferroni correction for multiple comparisons are shown (i.e. *p <* 0.05*/*15 = 0.0033); the remaining non-significant (n.s.) correlations are greyed out. Four exemplar significant relationships, indicated by dashed grey boxes in the lower triangular part of matrix A, are plotted in part B). For a supplementary analysis demonstrating differences in reliability and correspondence between the top and bottom 25% regions by volume (i.e. *<* Q_1_ VS *>* Q_3_), see supplementary Figure S3. (All reliability and correspondence results stem from T_2_w MRR scans.)

To help further illustrate the relationship of (some) regional statistics to average volume, we inspected the differences between the top and bottom 25% regions by volume (i.e. *<* Q_1_ VS *>* Q_3_) in all five reliability and correspondence statistics using Mann-Whitney U tests. For results, see supplementary Figure S3.

### Reliability and correspondence across contrasts and 64 mT resolutions

So far, we focused on results obtained using 64 mT T_2_w MRR scans, generated from three orthogonal scans using multi-resolution registration (Deoni et al., 2022), which showed the highest median between-scanner reliability and correspondence to 3 T 1 mm^3^ isotropic counterparts. Below, we summarise 64 mT reliability and correspondence to 3 T scans across all five 64 mT resolutions, for both T_1_w and T_2_w scans (Table 1). We focus on median values, and first and third quartiles (i.e. Md [Q_1_, Q^3^] across the 98 regional values of all five main reliability and correspondence statistics. Differences across scan resolutions were broadly consistent between T_2_w scans and were related to differences in voxel resolution and/or acquisition time. MRR scans, with the highest effective resolution, showed the highest performance across most measures, followed by high performance on (T_1_w) ICC and (T_2_w) CCC for ISO scans directly acquired at (lower) isotropic resolutions, with shorter thick-slice AXI, COR and SAG scans showing lower performance. Differences between the latter three acquisitions, as well as ISO scans on measures of Dice spatial overlap, were generally relatively minor. Differences across all five resolutions were more pronounced for cross-field correspondence than for 64 mT reliability statistics, and most prominent within measures of exact agreement across scanner field strengths (i.e., Lin’s CCC and median Dice_cor_), where MRR scans showed highest gains in performance relative to other resolutions (Table 1).

Furthermore, T_2_w scans showed moderately higher performance than T_1_w scans, possibly due to a slightly higher voxel resolution (e.g., T_2_w MRR scans have an effective resolution of 1.5 mm^3^ isotropic, compared to 1.6 mm^3^ isotropic for T_1_w MRR scans) as well as potential differences in partial volume artifacts at the time of data acquisition (see Discussion for details). Differences in performance across contrasts were again more prominent for measures of agreement across field strengths (Lin’s CCC and median Dice_cor_), where T_2_w scans showed highest gains in performance compared to T_1_w scans (e.g. with median performance gains of T_2_w MRR relative to T_1_w MRR scans of ΔCCC = 0.29, ΔDice_cor_ = 0.13, compared to ΔICC = 0.01, ΔDice_rel_ = 0.02 and Δr = 0.01).

### Measuring age-related changes in tissue volume

We wished to assess the extent to which 64 mT scans can be used to track changes in tissue volume, using linear models of both global and local volumes as a function of age, covarying for sex. Although our sample is relatively small and cross-sectional, the uniform distribution of participants across age (supplementary Figure S1) combined with marked changes in tissue volume across the lifespan (Bethlehem et al., 2022) result in clear age-related changes in both 3 T scans (Fig. 6A) and 64 mT scans (Fig. 6B). As expected, at the global level both field strengths show decreases in GM and WM volumes, and increases in CSF volumes, with similar effect sizes, albeit of reduced magnitude at 64 mT (including t-statistics for the effect of age, corresponding p-values and r^2^ for the variance in tissue volume explained by age and sex; Fig. 6 left).

**Figure 6:**
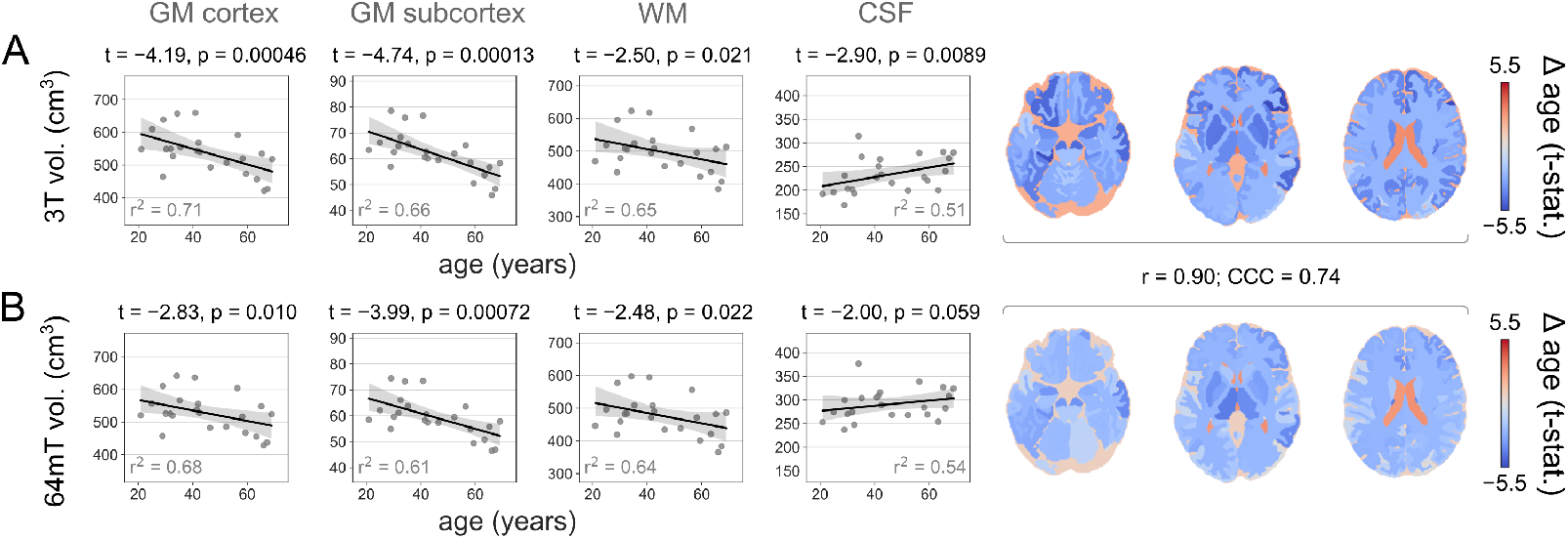
Changes in tissue volume from 3 T and 64 mT scans as a function of age. Rates of change of tissue volume as a function of age, in T_2_w A) 3 T scans and B) 64 mT MRR scans, across both four aggregate tissue classes (left) and individual regions (right). Both scanners show similar changes (values correspond to Pearson’s r and Lin’s CCC). For results across contrasts and 64 mT resolutions, see supplementary Table S3.

Local rates of change also showed excellent alignment across field strengths, with correspondence between local t-statistic maps of r = 0.90 and CCC = 0.74 (Fig. 6 right). Alignment was similar between local maps of the proportion of variance in tissue volumes explained by both age and sex (i.e., r^2^), which showed r = 0.78 and CCC = 0.71 between field strengths and across regions.

For correspondence of regional age-related changes in tissue volume across contrasts and 64 mT resolutions, see supplementary Table S3.

### Potential clinical applications

Finally, although our recruitment was restricted to neurologically healthy participants, we wished to explore the potential of clinical applications of ultra-low-field MRI in two distinct proof-of-concept analyses. For these analyses, we focused on T_2_w MRR ultra-low-field scans, which previously showed highest reliability and correspondence to high-field imaging.

In Fig. 7A, in a first hypothesis-driven clinically-relevant example, we highlight the between-scanner reliability, and correspondence between 64 mT and 3 T, of hippocampal volume, a key region of interest for the diagnosis and monitoring of Alzheimer’s Disease and other neurodegenerative processes. Reliability and correspondence statistics were derived as reported above. Similarly to other global and local tissue volumes, hippocampal volumes show high 64 mT test-retest reliability, and correspondence to 3 T (Fig. 7A).

**Figure 7:**
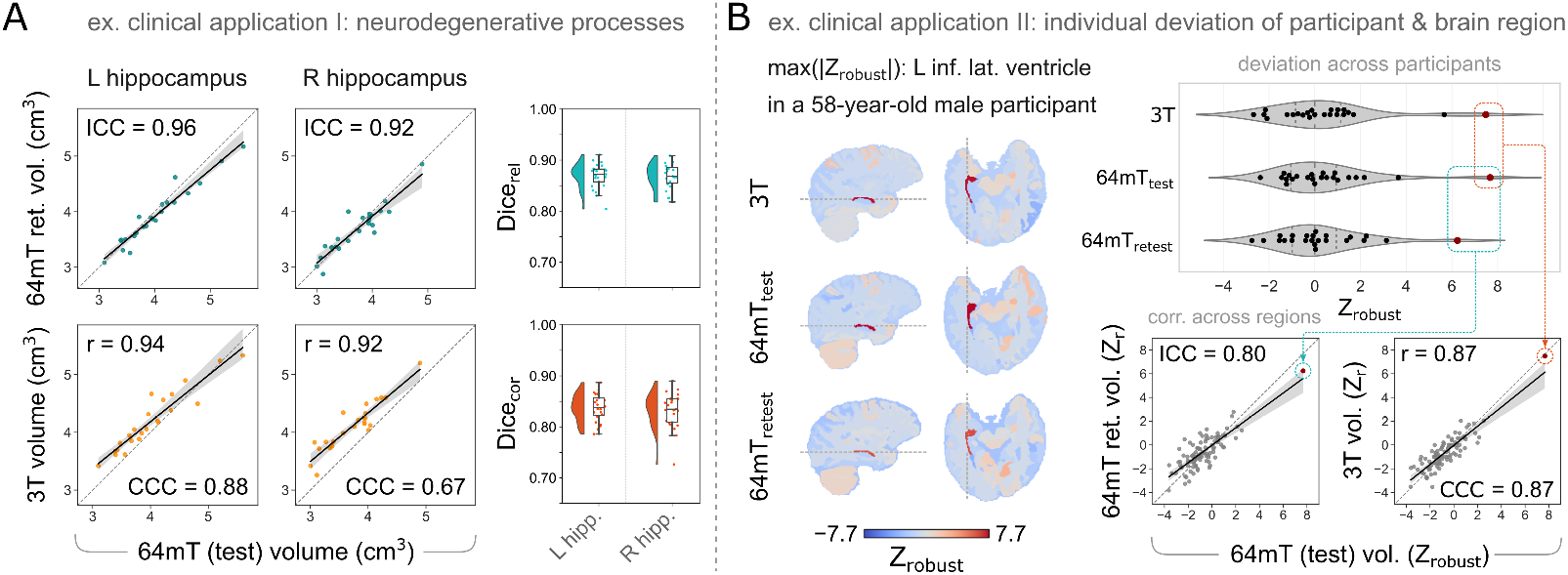
Two distinct example clinical applications of ultra-low-field morphometry. A) Example clinical application I: Hypothesis-driven focus on a region of interest for neurodegeneration. Hippocampal volume and segmentation are relevant for the diagnosis and monitoring of Alzheimer’s disease and other neurodegenerative processes, and show high 64 mT reliability (top) and correspondence to 3 T MRI (bottom), both in volume correlations (left) and Dice overlap of segmentations (right). B) Example clinical application II: Data-driven identification of individual deviation from the norm. Individual robust Z-scores (Z_*robust*_) of tissue volumes (corrected for age and sex) highlight the same outlier individual, and region, across scanners.

Additionally, in a second data-driven clinically-relevant example, we investigated the potential of 64 mT scans to identify individuals who deviate from the norm (Marquand et al., 2019). We first corrected all local tissue volumes for the effects of age and sex by retaining the residuals of linear models (described above), and subsequently computed non-parametric equivalents of the Z-score by robust-scaling corrected volumes, within each region, across participants (leading to a robust Z-score, or Z_*robust*_) (Leys et al., 2013). We then identified the participant and region showing the greatest deviation from the median in T_2_w high-field scans, and visualized both (i) the corrected volume of this region from all three scanners, across participants, and (ii) the reliability and correspondence of all regional deviations in this participant (Fig. 7B). The same participant - a 58-year-old-male - was identified as the greatest outlier across scanners. Even after correcting regional volume for age and sex, regional deviations within this participant showed high reliability and correspondence across regions.

## Discussion

We reported results from 23 healthy adult participants, scanned on two identical portable 64 mT ultra-low-field MRI scanners, and a 3 T high-field scanner. We segmented T_1_w and T_2_w scans acquired at a range of resolutions into 98 structures across 4 tissue types and estimated their volumes. We showed excellent test-retest reliability of morphometric measurements extracted from 64 mT scans, and high correspondence to 3 T MRI, using volume correlations and Dice overlap of both global tissue classes and individual regions. Reliability and correspondence were highest using T_2_w MRR scans, obtained using multi-resolution registration of three orthogonal 64 mT acquisitions with thick slices but high in-plane resolution; however, these lower-resolution scans also showed excellent performance individually, as did scans directly acquired at (reduced) isotropic resolution. In additional analyses, we showcased the potential of ultra-low-field imaging for assessment of age-related changes in tissue volume, as well as potential clinical applications.

### Determinants of reliability and correspondence

We reported higher between-scanner 64 mT reliability and correspondence to 3 T scans for morphometric measures extracted from T_2_w ultra-low-field scans, than from T_1_w scans, in this dataset. This may be due to several factors. First, the T_2_w scans were of slightly higher resolution across acquisitions, including for example 1.5×1.5 mm^2^ within the high-resolution plane for AXI, COR and SAG scans, compared to 1.6×1.6 mm^2^ for corresponding T_1_w scans; or a resulting effective resolution of 1.5 mm^3^ isotropic for T_2_w MRR scans, compared to 1.6 mm^3^ isotropic for T_1_w MRR scans (see supplementary Table S1 for details). Additionally, within the Hyperfine Swoop software version used at the time of acquisition (rc8.6.0), 64 mT T_1_w sequence parameters resulted in non-zero CSF signal and partial-volume signal voids at tissue interfaces, possibly explaining lower correspondence to highfield imaging. Updated parameters in more recent software versions now yield T_1_w scans with nulled CSF signal whilst retaining grey/white matter contrast, which may result in greater correspondence to high-field scans. We note also that partial volume effects may be responsible for substantial overestimation of CSF volume, in both T_1_w and T_2_w. Similar overestimates have been reported by other groups Hsu et al. (2025).

Both reliability and correspondence to 3 T were higher for 64 mT acquisitions with higher (effective) resolution; namely, highest for 1.5 mm^3^ (and 1.6 mm^3^) isotropic MRR scans, followed by isotropic ISO scans and non-isotropic AXI / COR / SAG scans, all with identical voxel volumes (within contrasts). Differences between these lower-resolution scans were relatively minor, with a tendency for COR scans (with higher resolution within the coronal plane) to show higher performance. These results are in line with prior work showing that morphometric reliability is affected by voxel size and geometry, with larger voxels leading to lower morphometric reliability and accuracy, in both high-field (Wonderlick et al., 2009; Haller et al., 2016; Mulder et al., 2019) and ultra-low-field scans (Sorby-Adams et al., 2024). While the SynthSeg+ algorithm used here for segmentation and volume estimation is designed to run on scans of any contrast and resolution (Billot et al., 2023), its performance remains partially dependent on variations in image properties.

We also found significant relationships of some reliability and correspondence statistics to each other, and to region size (as estimated using high-field MRI). We note that only some - not all - reliability and correspondence statistics were related to volume, and to each other, across regions. Still, we suggest that these relationships can be summarized as: “the bigger (the region), the better (the agreement)”. This aligns with prior research demonstrating a tendency for smaller (sub-)regions to have lower morphometric reliability (Morey et al., 2010; Whelan et al., 2016; Madan and Kensinger, 2017; Elliott et al., 2023).

Most prior work on the correspondence of 64 mT morphometric measurements to 3 T (Iglesias et al., 2023; Cooper et al., 2024; Sorby-Adams et al., 2024; Islam et al., 2025; Pretzsch et al., 2025) relied on the use of pre-trained deep-learning algorithms to enhance the quality of ultra-low-field scans (Iglesias et al., 2021). Here we show excellent reliability and correspondence across fields without prior application of image quality transfer (IQT), similarly to recent work (Hsu et al., 2025) largely focused on coarse tissue types. This eliminates the potential for artifacts introduced by neural networks, which may subtly alter anatomy while super-resolving images (Zhao et al., 2020; Lin et al., 2023), particularly if the input image deviates from the training dataset; for example, if applying a super-resolution model trained on healthy adults to paediatric and/or clinical scans (Iglesias et al., 2021, 2023; Baljer et al., 2024). Still, IQT may be useful for many applications, including qualitative visual assessment of scans for clinical purposes, or application of pre-processing tools which may require images of a specific contrast and/or resolution (e.g., “standard” 1 mm^3^ isotropic T_1_w scans; (Iglesias et al., 2021)). We also note that the Hyperfine Swoop 64 mT system includes a deep-learning processing step to reduce image blurring and noise that is performed on the scanner during image reconstruction (Krupinski et al., 2023); as such, this impacts all scans acquired using the Hyperfine Swoop ultra-low-field system.

Our results may help inform the choice of protocol for future 64 mT studies acquiring T_1_w and/or T_2_w scans. The specific choice of sequences is likely to depend on the question of interest, and time available to scan each participant; however, based on our results, for the purposes of segmentation and tissue volume estimation, it seems sensible to prioritize thick-slice T_2_w scans acquired in three orthogonal orientations, which can be combined into a single isotropic scan using multi-resolution registration (Deoni et al., 2022). Beyond the analytic applications explored here, investigators can use the publicly available data to validate and optimize alternative or custom analytic pipelines, to maximize the reliability and/or cross-field correspondence of desired imaging phenotypes.

### Ultra-low-field MR imaging

Analysis of MR images for morphometric measurements relies on the ability to discern anatomical structures based on the MR image contrast. In brain imaging, white matter, gray matter, and CSF can be clearly distinguished as their T_1_, T_2_, and T_2_* relaxation times are different. These differences are driven by the underlying tissue microstructure, e.g., myelin, iron, and water content. However, these relaxation times change with Larmor frequency thus making the tissue contrast, i.e., the basis for our morphometric measurements, dependent on the field strength of the MR system.

The field strength dependence of relaxation times is complicated, non-linear, and depends on numerous factors, but it generally holds that T_1_ is shorter at lower field strength (Rooney et al., 2007), T_2_ only shows small changes, if any (Stanisz et al., 2005; Jordanova et al., 2023), and T_2_* increases (Peters et al., 2007). Since the relaxation times, which produce the tissue contrast, do not change in the same way with field strength, different pulse sequence parameters are required at high and low field strengths (see supplementary Table S1), making it non-trivial to produce identical image contrast. This could impact morphometric measurements if brain regions are heterogeneous in their tissue composition (e.g., Fukunaga et al., 2010).

Another factor to consider is the signal-to-noise ratio (SNR) which is inherently tied to the field strength, resulting in greatly reduced SNR at 64 mT compared to 3 T (Marques et al., 2019). To partly compensate for this, ultra-low-field MR systems use large voxel size (here, 1.5×1.5×5 mm^3^ compared to 1×1×1 mm^3^ for T_2_w 64 mT and 3 T product sequences), and more signal averaging, resulting in longer acquisition times. When participants lie in the scanner for a longer period of time, the risk of motion artifacts also increases, an issue that could be particularly relevant for older and paediatric populations (Madan, 2018; Padormo et al., 2023).

The ultra-low-field MRI system used in this study, Hyperfine Swoop, is one of the few commercially available systems in the rapidly growing space of ultra-low-field MRI. With reduced costs and more lenient siting requirements, ultra-low-field systems are being developed by research groups around the world with various designs for both brain (O’Reilly et al., 2019; McDaniel et al., 2019; He et al., 2020; Liu et al., 2021; Cooley et al., 2021; Guallart-Naval et al., 2022) and whole-body imaging (Zhao et al., 2024). A unique aspect of ultra-low-field MR systems is the ability to make them portable and able to operate in a normal room, by the bedside in a clinical setting (Mazurek et al., 2021; Balaji et al., 2024), in resource-constrained environments (Chetcuti et al., 2022; Abate et al., 2024) or even in an ambulance during active transport (Roberts et al., 2023). However, this also means that the environment can change from scan to scan, including both temperature and electromagnetic interference. The latter is, in particular, an issue that is unique to portable MR systems and a growing field of research. The fact that the scanner may be located in a non-controlled environment should be considered as a reproducibility aspect in future studies.

### Limitations and further work

We only investigated reliability and correspondence of tissue volumes and underlying segmentations, obtained using a single pre-processing algorithm (SynthSeg+; Billot et al., 2023). A much wider range of pre-processing and analysis approaches could be applied to systematically explore their impact on the reliability and correspondence of derived imaging phenotypes (e.g., Heinen et al., 2016; Sederevičius et al., 2021). However, raw images used in this study are publicly available, and we envisage that they will constitute a useful resource for researchers interested in developing and validating other pre-processing and analysis approaches for 64 mT MRI.

We note that the MRR command was initially developed for the construction of group templates from a large sample of participants, and that its application to a relatively small number of (ultralow-field MRI) scans acquired in orthogonal planes may lead to visual artifacts such as cross-hatching. While this approach retains favourable performance in our use-case, ROI-based morphometry, despite such artifacts, this may not be the case in voxel-wise approaches such as voxel-based morphometry (VBM). Further exploration of MRR, including in particular more systematic investigation of its numerous pre-processing steps and parameters, may lead to further recommendations regarding this approach. In the meantime, results presented here and elsewhere (e.g., Deoni et al., 2022; Briski et al., 2024; Ringshaw et al., 2025) indicate the promise of this method.

Additionally, our data and conclusions are limited to healthy adult participants, and additionally tempered by a modest sample size. Acquisition and open sharing or similar paired high-and ultra-low-field scans from paediatric and/or clinical populations, as well as similar healthy adult scans, will help accelerate development and validation of analytic approaches for ultra-low-field imaging. For example, larger datasets could help establish “confidence intervals” and re-calibration values for ultra-low-field morphometric estimates, relative to high-field counterparts. Still, exact correspondence to 3 T reference measurements may not be required for many applications of 64 mT morphometry, including for example longitudinal assessment of neurodevelopmental and/or clinical change within individuals, where measures at new time-points can be referenced to prior scans.

We note that our data are generally of very high quality, thanks in part to a focus on healthy adult participants who can lie still in the scanner; correspondence is likely to be lower within paediatric, elderly and/or clinical populations, who are likely to exhibit increased head motion (Madan, 2018; Padormo et al., 2023). Additionally, we used image processing tools developed for healthy adult participants (i.e., SynthSeg+; Billot et al., 2023), which may not perform as well on alternative populations. However, these limitations may be offset by developments in ultra-low-field imaging hardware and software, as well as software used for image enhancement and pre-processing.

Our results indicate greater ultra-low-field reliability and correspondence to high-field MRI of tissue volumes, than of the underlying regional segmentations. This is unsurprising, as local errors in the labeling of individual voxels are compounded when quantifying the overlap of segmentation labels, but may cancel out when calculating an aggregate measure of volume. For example, a structure may be assigned the same number of voxels in two scans, but these voxels will not necessarily overlap between scans (Kahhale et al., 2023). Going forward, for a more systematic assessment of the spatial overlap of segmentations, additional measures may be useful (Reinke et al., 2024). We also note that 3 T segmentations and corresponding volumes were used as reference, although these may show imperfections as well. While all segmentations were visually quality-controlled, further work on the correspondence of morphometric measures across field strengths would benefit from paired high-and ultra-low-field data with manually-labeled, “gold-standard” segmentations (e.g., Klein and Tourville, 2012).

The current study has only investigated anatomical imaging sequences as these are currently the most widely used at ultra-low field. The low SNR makes it difficult to utilize techniques which require rapid imaging, such as perfusion or functional imaging. Some image contrasts are difficult to achieve simply due to the MR physics. BOLD contrast, as used in functional MR, currently rely on contrast change due to susceptibility changes in oxygenated blood, an effect which is vanishingly small at ultra-low field. Functional imaging studies at ultra-low field strength would therefore have to consider alternative methods of probing brain function. We note that the Hyperfine Swoop 64 mT scanner can currently be used to acquire limited diffusion MRI data, including a single-orientation diffusion-weighted volume and an apparent diffusion coefficient (ADC) map; reliability and of ultra-low-field diffusion MRI, and correspondence to 3 T, is an avenue for future work.

Another aspect for wider applicability is the anatomical coverage of ultra-low-field systems. Many are currently built for a specific region, such as the brain (McDaniel et al., 2019; O’Reilly et al., 2019; He et al., 2020; Cooley et al., 2021; Liu et al., 2021; Kimberly et al., 2023) or extremities (Guallart-Naval et al., 2022), as this can make the system more affordable and may result in optimal performance for a given application. However, recent advancements have shown the possibility of performing whole body imaging on a 50 mT MR system (Zhao et al., 2024).

## Conclusion

Volume estimates from portable ultra-low-field MRI scans show excellent test-retest reliability and correspondence to high-field counterparts. Reliability and correspondence to 3 T were highest for T_2_w 64 mT scans reconstructed using MRR. Our results pave the way for clinical applications of ultra-low-field imaging morphometry, such as modeling of individual deviations from the norm, across development and disease. In particular, the excellent between-scanner reliability of 64 mT morphometry and high correspondence to 3 T augurs well for the pooling of ultra-low-field MRI data across sites to help address issues of brain health at scale across the globe.

## Data availability

All raw imaging data used in this manuscript are available at https://openneuro.org/datasets/ds006557/versions/1.0.0. Pre-processed derivatives will be made available upon publication.

## Code availability

All code used for pre-processing and statistical analysis will be made available upon publication.

## Author contributions

F.V., C.B., R.L., S.C.R.W. designed the study. F.P., T.C.W., D.J.W., F.D.A., E.L. implemented MRI sequences and provided MRI technical support. F.V., C.B., P.C., T.A. acquired the data. F.V., N.J.B., L.B. pre-processed the data. F.V., U.B. performed quality control of the data. F.V. performed analyses. T.C.B., A.V., S.C.L.D., R.J.M. provided conceptual support and advice. F.V., C.B., E.L. wrote the manuscript. All authors discussed results and commented on the manuscript.

## Acknowledgements

This study was supported by a National Institute for Health and Care Research (NIHR) Biomedical Research Centre Early Career Investigator Award to F.V., and by the Bill and Melinda Gates Foundation UNITY project [INV-032788; INV-047888]. T.A. receives funding from the MRC via a Senior Clinical Fellowship [MR/Y009665/1] and the MRC Centre for Neurodevelopmental Disorders, King’s College London [MR/N026063/1]. A.V.V. is funded by the National Institute for Health and Care Research as NIHR Clinical Lecturer and supported by the NIHR Maudsley Biomedical Research Centre and NIHR HealthTech Research Centre in Brain Health at King’s College London and South London and Maudsley NHS Foundation Trust. S.C.R.W. is funded by the NIHR Biomedical Research and HealthTech Centres at the South London & Maudsley Hospital.

