## Supplementary Information for "Ultra-low-field brain MRI morphometry: test-retest reliability and correspondence to high-field MRI"

### Supplementary Text

#### Multi-Resolution Registration

We used multi-resolution registration (Deoni et al., 2022) to combine the three orthogonal non-isotropic 64 mT scans (AXI, COR, SAG) into a single “MRR” scan, of a higher effective resolution, using ANTs (Avants et al., 2009).

We first pre-aligned scans, and resampled them to $1\times1\times1$ mm$^3$, by rigid-registering them to the high-field scan of the corresponding contrast (i.e. $T_2$w or $T_1$w), using *antsRegistrationSyN.sh*.

We subsequently applied the *antsMultivariateTemplateConstruction2.sh* command, with the following options (based on Deoni et al., 2022):

- Transformation model: “SyN” (Greedy SyN)
- Similarity metric used for pairwise registration: “MI” (mutual information)
- Maximum iteration limit: 12

Other options were left unchanged from default values, including:

- Image statistic used to summarize images: mean of normalized intensities (As described in the aforementioned script: *“Normalization here means dividing each image by its mean intensity.”)*
- Sharpening applied to template at each iteration: Laplacian
- Gradient step size: 0.25

For further details, see the accompanying code (the link is located within the main manuscript).

#### Supplementary Figures

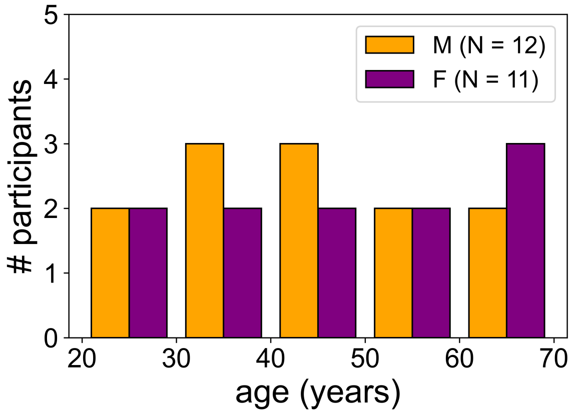

**Figure S1**: **Participant demographics.** Distribution of participants as a function of age, stratified by sex.

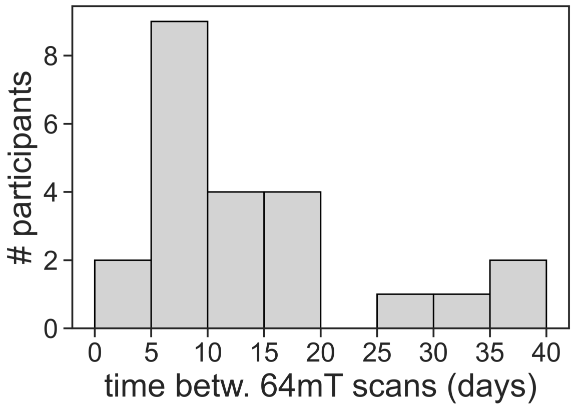

**Figure S2**: **Time between 64 mT scans, within participants.** The first 64 mT scan was acquired on the same day as, and immediately following, the 3 T scan.

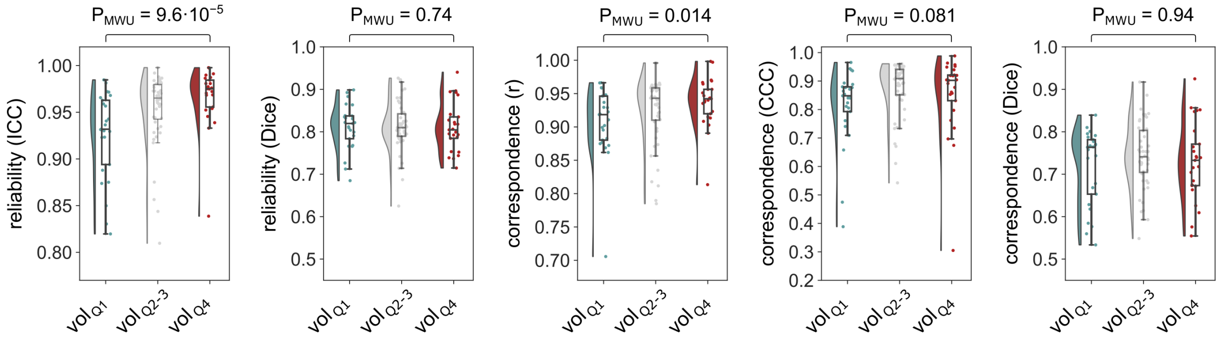

**Figure S3**: **Differences in reliability and correspondence between small and large regions.** The P-value ($P_{MWU}$) corresponds to a Mann-Whitney U test between the bottom and top quartile by average region volume (extracted from high-field scans), for each reliability and correspondence statistic. The middle two quartiles are included for illustration purposes.

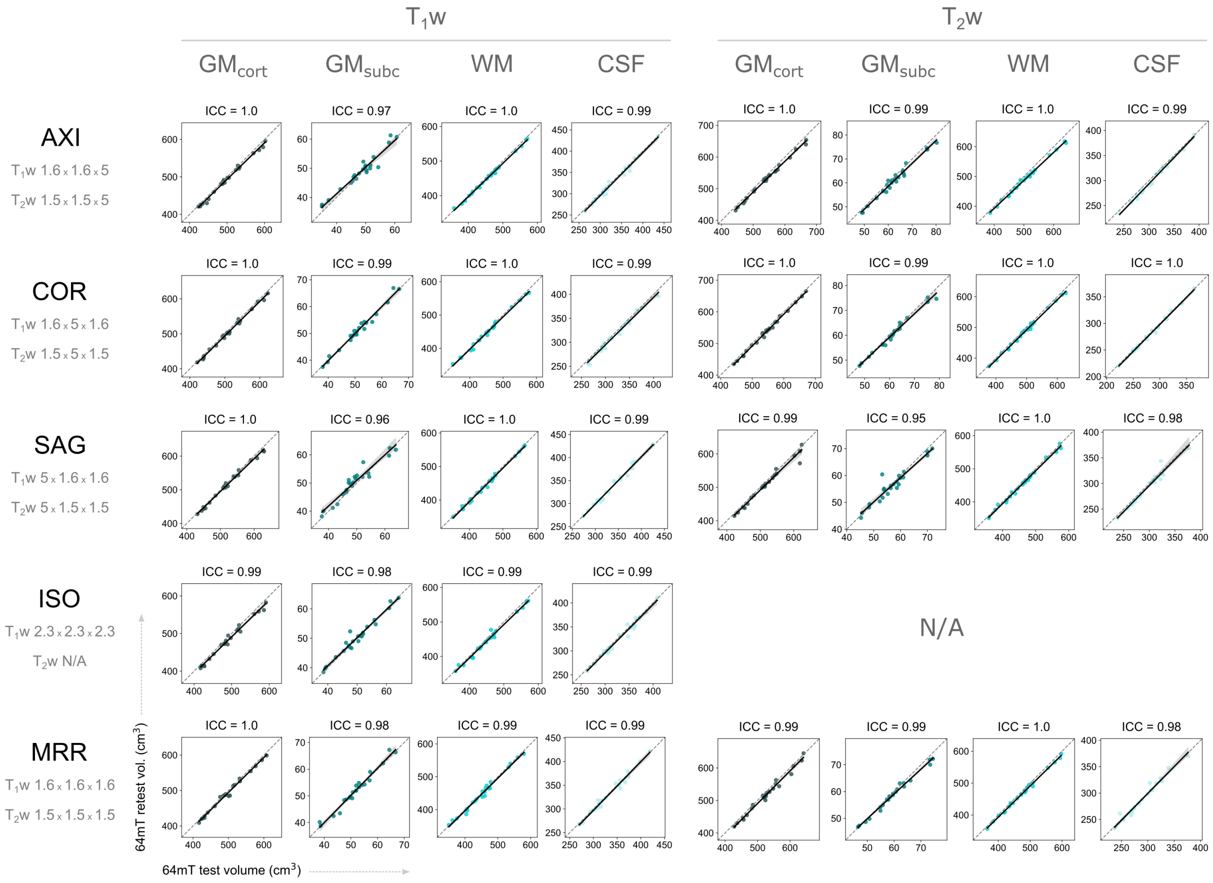

**Figure S4**: **Global reliability of 64 mT tissue volumes, across contrasts and resolutions.**

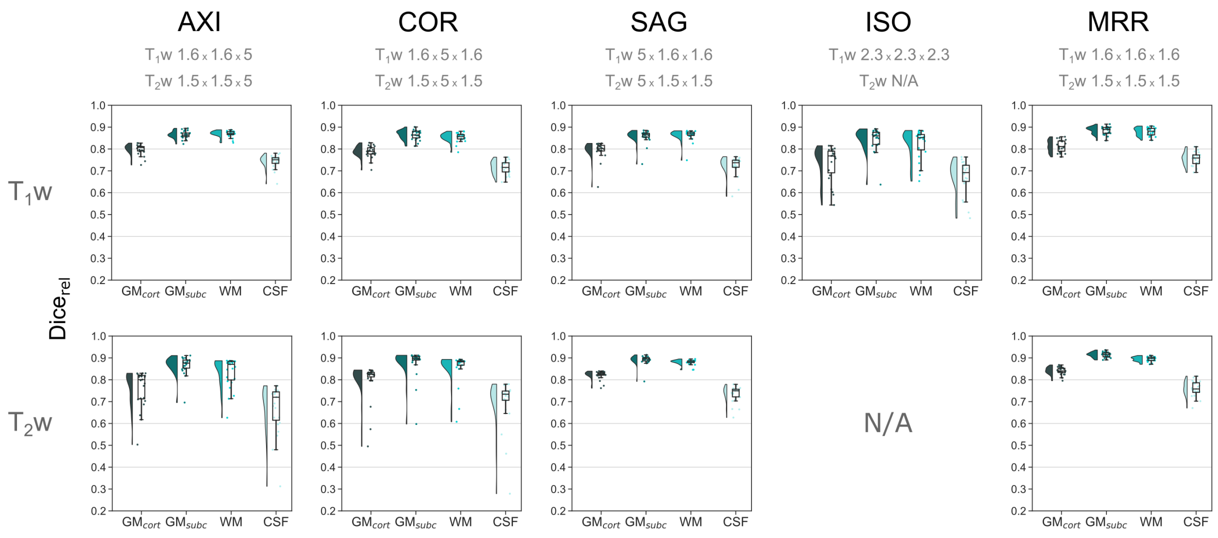

**Figure S5**: **Global reliability of 64 mT segmentations, across contrasts and resolutions.**

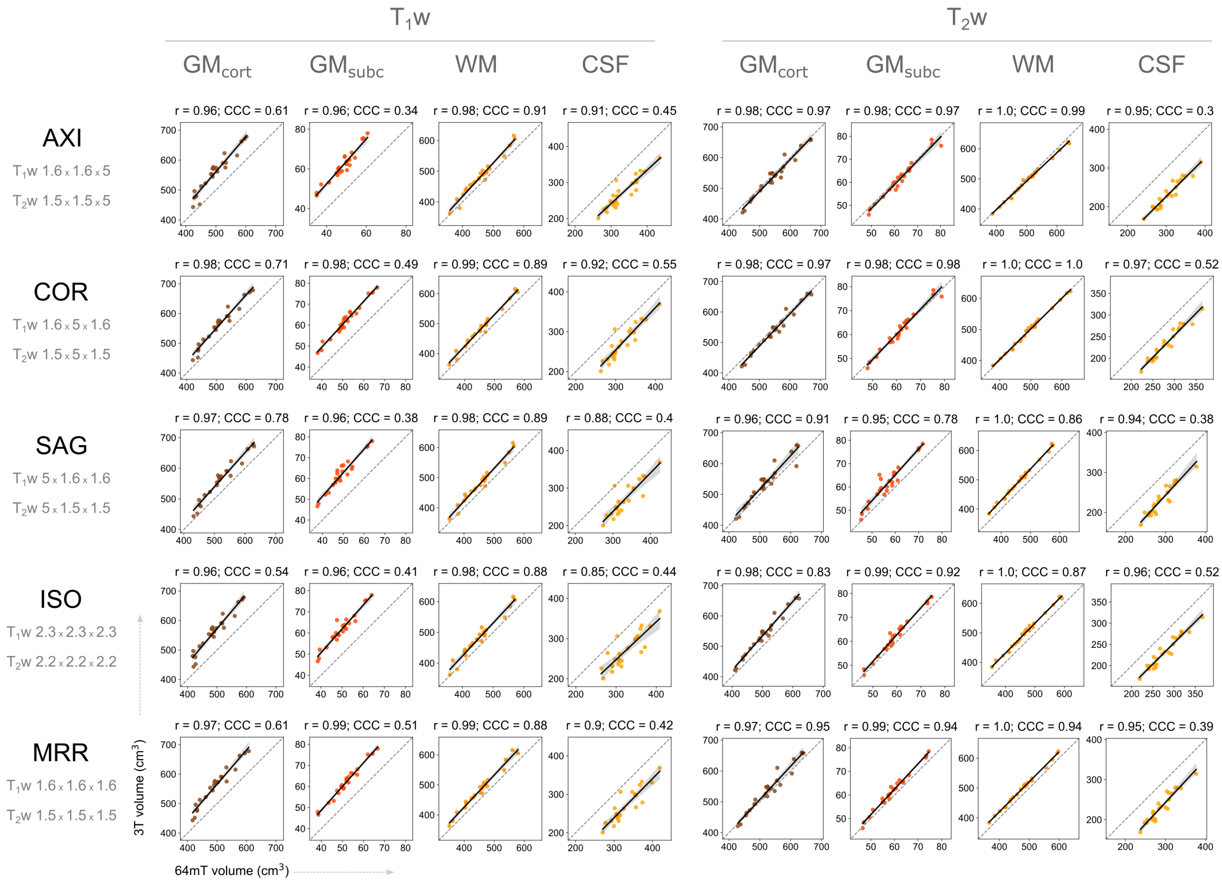

**Figure S6**: **Global correspondence of 64 mT tissue volumes to 3 T, across contrasts and resolutions.**

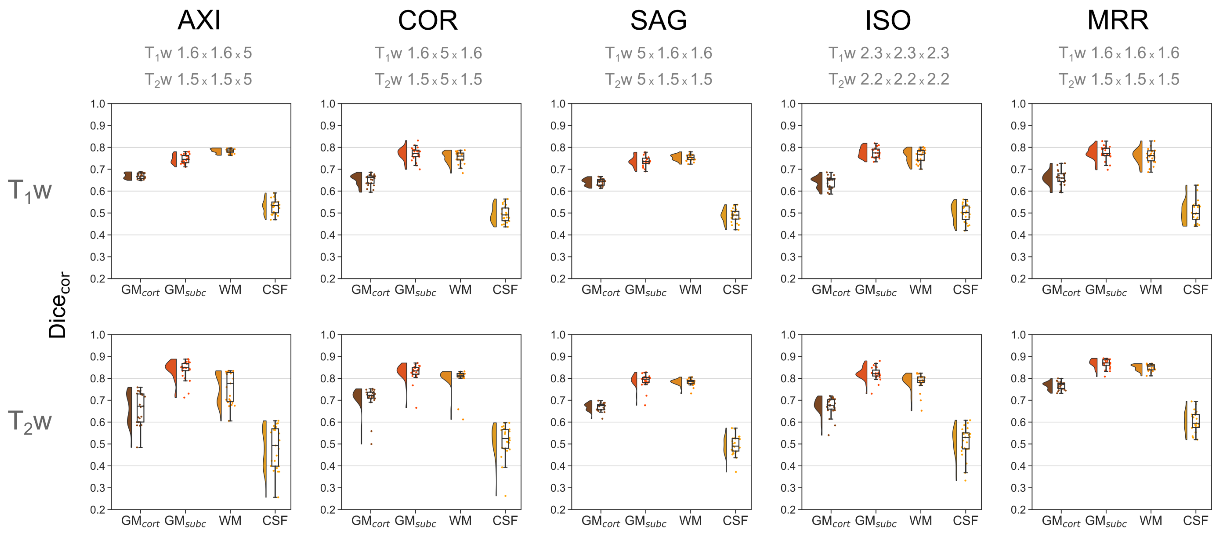

**Figure S7: Global correspondence of 64 mT segmentations to 3 T, across contrasts and resolutions.**

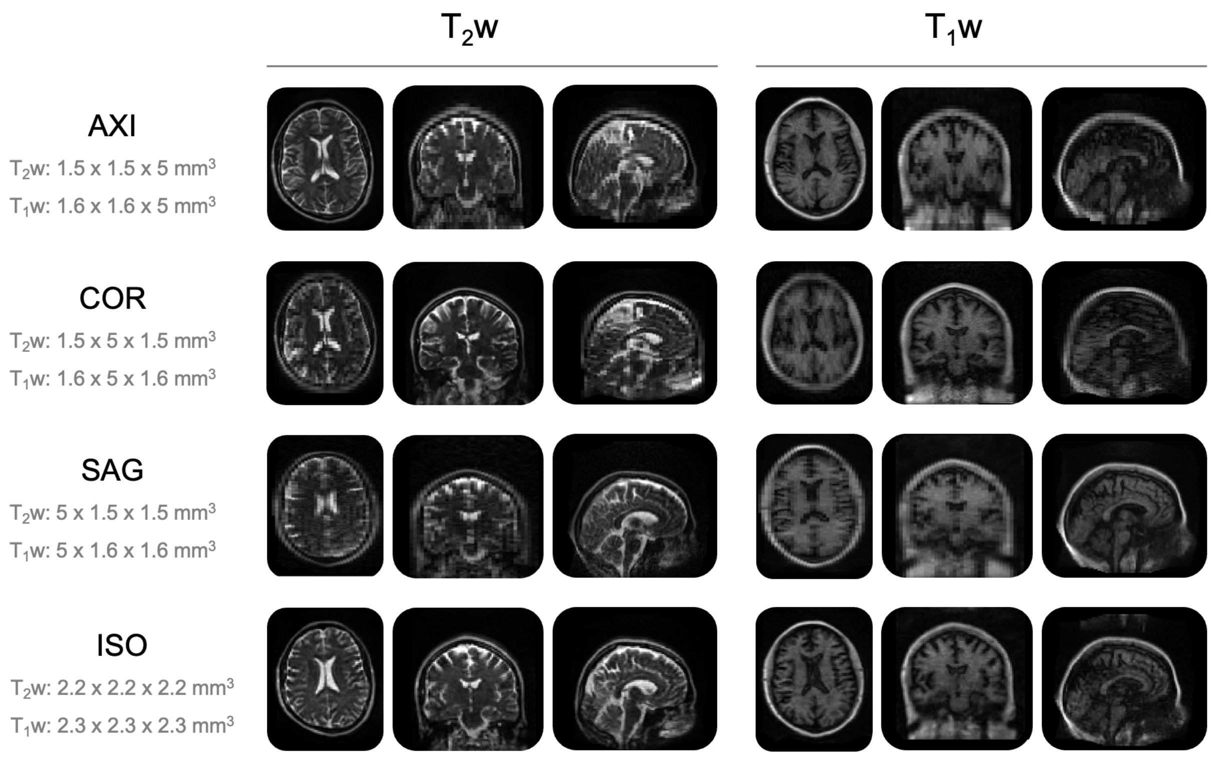

**Figure S8: All raw 64 mT scans in all three planes for one example participant.**

### Supplementary Tables

**Table S1**: **Acquisition parameters for 64 mT and 3 T scans.** For all 64 mT scans, the FOV is 22×18×20 cm. *The MRR volume consists of a "multi-resolution registration" reconstruction (Deoni et al., 2022) of the AXI, COR and SAG volumes. †Accordingly, the MRR acquisition time corresponds to the sum of the acquisition times of the AXI, COR and SAG volumes (i.e. it does not require additional scan time beyond these three scans).

| field | contrast | name | resolution (mm) | vox. vol. ($mm^3$) | matrix size | TR / TE / TI (s) | duration (min:s) |
|---|---|---|---|---|---|---|---|
| 64 mT | $T_1$w | AXI | 1.6×1.6×5 | 12.8 | 112×138×40 | 1.5 / 0.007 / 0.3 | 6:16 |
| | | COR | 1.6×5×1.6 | 12.8 | 112×44×124 | 1.5 / 0.007 / 0.3 | 5:31 |
| | | SAG | 5×1.6×1.6 | 12.8 | 36×138×125 | 1.5 / 0.007 / 0.3 | 6:13 |
| | | ISO | 2.3×2.3×2.3 | 12.8 | 78×94×86 | 1.5 / 0.006 / 0.3 | 9:26 |
| | | MRR* | 1.6×1.6×1.6 | 4.1 | N/A | N/A | 18:00† |
| | $T_2$w | AXI | 1.5×1.5×5 | 11.3 | 120×146×40 | 2.0 / 0.17 / N/A | 6:25 |
| | | COR | 1.5×5×1.5 | 11.3 | 120×40×134 | 2.0 / 0.19 / N/A | 5:21 |
| | | SAG | 5×1.5×1.5 | 11.3 | 36×146×146 | 2.0 / 0.20 / N/A | 5:49 |
| | | ISO | 2.2×2.2×2.3 | 11.4 | 80×96×92 | 2.0 / 0.15 / N/A | 9:49 |
| | | MRR* | 1.5×1.5×1.5 | 3.4 | N/A | N/A | 17:35† |
| 3 T | $T_1$w | MPRAGE | 1.0×1.0×1.0 | 1.0 | 208×256×256 | 2.5 / 0.003 / 1.06 | 5:01 |
| | $T_2$w | FSE | 1.0×1.0×1.0 | 1.0 | 208×256×256 | 3.2 / 0.07 / N/A | 6:06 |

**Table S2**: **Summary of percentage difference in tissue volumes from 64 mT scans relative to 3 T scans, across contrasts and resolutions.** Global values correspond to the median across 4 global tissue classes; local values correspond to the median, and first and third quartiles, across 98 structures segmented by SynthSeg+.

| | contrast | MRR<br>$T_1$w 1.6×1.6×1.6<br>$T_2$w 1.5×1.5×1.5 | ISO<br>$T_1$w 2.3×2.3×2.3<br>$T_2$w 2.2×2.2×2.2 | AXI<br>$T_1$w 1.6×1.6×5<br>$T_2$w 1.5×1.5×5 | COR<br>$T_1$w 1.6×5×1.6<br>$T_2$w 1.5×5×1.5 | SAG<br>$T_1$w 5×1.6×1.6<br>$T_2$w 5×1.5×1.5 |
|---|---|---|---|---|---|---|
| **Global** (Md) | $T_1$w | −9.3 | −9.5 | −8.2 | −7.7 | −7.0 |
| | $T_2$w | −2.6 | −5.3 | +1.6 | +0.8 | −4.9 |
| **Local** (Md [$Q_1$,$Q_3$]) | $T_1$w | −11.3 [−15.4,−7.3] | −12.7 [−22.4,−7.4] | −12.3 [−23.1,−5.9] | −11.7 [−20.5,−4.0] | −10.1 [−21.5,−3.0] |
| | $T_2$w | −1.6 [−4.4,+0.9] | −3.8 [−11.0,−0.5] | +2.3 [−5.7,+7.1] | +2.6 [−6.4,+7.4] | −3.7 [−9.8,+1.6] |

**Table S3**: **Summary of correspondence between local age-statistics extracted from 64 mT and 3 T scans, across contrasts and resolutions.** Linear models of regional tissue volume as a function of age and sex were used to extract t-statistics for the effect of age, and variance explained by age and sex ($r^2$), separately for 3 T and 64 mT scans. The correspondence of both effect sizes between field strengths, across regions was quantified using Pearson's r, and Lin's CCC, across both contrasts and all 64 mT scan resolutions.

| effect size | corresp. statistic | contrast | MRR<br>$T_1$w 1.6×1.6×1.6<br>$T_2$w 1.5×1.5×1.5 | ISO<br>$T_1$w 2.3×2.3×2.3<br>$T_2$w 2.2×2.2×2.2 | AXI<br>$T_1$w 1.6×1.6×5<br>$T_2$w 1.5×1.5×5 | COR<br>$T_1$w 1.6×5×1.6<br>$T_2$w 1.5×5×1.5 | SAG<br>$T_1$w 5×1.6×1.6<br>$T_2$w 5×1.5×1.5 |
|---|---|---|---|---|---|---|---|
| **t-stat. (age)** | **Pearson's r** | $T_1$w | 0.83 | 0.78 | 0.79 | 0.82 | 0.77 |
| | | $T_2$w | 0.90 | 0.84 | 0.83 | 0.84 | 0.77 |
| | **Lin's CCC** | $T_1$w | 0.70 | 0.61 | 0.60 | 0.68 | 0.53 |
| | | $T_2$w | 0.74 | 0.70 | 0.67 | 0.70 | 0.54 |
| $\mathbf{r^2}$ | **Pearson's r** | $T_1$w | 0.72 | 0.71 | 0.57 | 0.65 | 0.56 |
| | | $T_2$w | 0.78 | 0.70 | 0.63 | 0.68 | 0.54 |
| | **Lin's CCC** | $T_1$w | 0.69 | 0.54 | 0.39 | 0.49 | 0.37 |
| | | $T_2$w | 0.71 | 0.59 | 0.53 | 0.55 | 0.37 |

**Table S4**: **Assignment of SynthSeg+ regions to tissue classes.** The first column contains region names, as provided by SynthSeg+ (Billot et al., 2023). The second column contains the assignment of each region to one of four tissue classes; $\mathrm{GM}_{\mathrm{cort}}$: cortical gray matter, $\mathrm{GM}_{\mathrm{subc}}$: subcortical gray matter, WM: white matter, CSF: cerebrospinal fluid (cerebellar regions and the brainstem were not assigned to any of the four classes). The third colum contains the label of each region in the .nii.gz segmentation files generated by SynthSeg+.

| name | tissue | label |
|---|---|---|
| Left-Cerebral-White-Matter | WM | 2 |
| Left-Lateral-Ventricle | CSF | 4 |
| Left-Inf-Lat-Vent | CSF | 5 |
| Left-Cerebellum-White-Matter | - | 7 |
| Left-Cerebellum-Cortex | - | 8 |
| Left-Thalamus-Proper | $\mathrm{GM}_{\mathrm{subc}}$ | 10 |
| Left-Caudate | $\mathrm{GM}_{\mathrm{subc}}$ | 11 |
| Left-Putamen | $\mathrm{GM}_{\mathrm{subc}}$ | 12 |
| Left-Pallidum | $\mathrm{GM}_{\mathrm{subc}}$ | 13 |
| 3rd-Ventricle | CSF | 14 |
| 4th-Ventricle | CSF | 15 |
| Brain-Stem | - | 16 |
| Left-Hippocampus | $\mathrm{GM}_{\mathrm{subc}}$ | 17 |
| Left-Amygdala | $\mathrm{GM}_{\mathrm{subc}}$ | 18 |
| CSF | CSF | 24 |
| Left-Accumbens-area | $\mathrm{GM}_{\mathrm{subc}}$ | 26 |
| Left-VentralDC | $\mathrm{GM}_{\mathrm{subc}}$ | 28 |
| Right-Cerebral-White-Matter | WM | 41 |
| Right-Lateral-Ventricle | CSF | 43 |
| Right-Inf-Lat-Vent | CSF | 44 |
| Right-Cerebellum-White-Matter | - | 46 |
| Right-Cerebellum-Cortex | - | 47 |
| Right-Thalamus-Proper | $\mathrm{GM}_{\mathrm{subc}}$ | 49 |
| Right-Caudate | $\mathrm{GM}_{\mathrm{subc}}$ | 50 |
| Right-Putamen | $\mathrm{GM}_{\mathrm{subc}}$ | 51 |
| Right-Pallidum | $\mathrm{GM}_{\mathrm{subc}}$ | 52 |
| Right-Hippocampus | $\mathrm{GM}_{\mathrm{subc}}$ | 53 |
| Right-Amygdala | $\mathrm{GM}_{\mathrm{subc}}$ | 54 |
| Right-Accumbens-area | $\mathrm{GM}_{\mathrm{subc}}$ | 58 |
| Right-VentralDC | $\mathrm{GM}_{\mathrm{subc}}$ | 60 |
| ctx-lh-bankssts | $\mathrm{GM}_{\mathrm{cort}}$ | 1001 |
| ctx-lh-caudalanteriorcingulate | $\mathrm{GM}_{\mathrm{cort}}$ | 1002 |
| ctx-lh-caudalmiddlefrontal | $\mathrm{GM}_{\mathrm{cort}}$ | 1003 |
| ctx-lh-cuneus | $\mathrm{GM}_{\mathrm{cort}}$ | 1005 |
| ctx-lh-entorhinal | $\mathrm{GM}_{\mathrm{cort}}$ | 1006 |
| ctx-lh-fusiform | $\mathrm{GM}_{\mathrm{cort}}$ | 1007 |
| ctx-lh-inferiorparietal | $\mathrm{GM}_{\mathrm{cort}}$ | 1008 |
| ctx-lh-inferiortemporal | $\mathrm{GM}_{\mathrm{cort}}$ | 1009 |
| ctx-lh-isthmuscingulate | $\mathrm{GM}_{\mathrm{cort}}$ | 1010 |
| ctx-lh-lateraloccipital | $\mathrm{GM}_{\mathrm{cort}}$ | 1011 |
| ctx-lh-lateralorbitofrontal | $\mathrm{GM}_{\mathrm{cort}}$ | 1012 |
| ctx-lh-lingual | $\mathrm{GM}_{\mathrm{cort}}$ | 1013 |
| ctx-lh-medialorbitofrontal | $\mathrm{GM}_{\mathrm{cort}}$ | 1014 |
| ctx-lh-middletemporal | $\mathrm{GM}_{\mathrm{cort}}$ | 1015 |
| ctx-lh-parahippocampal | $\mathrm{GM}_{\mathrm{cort}}$ | 1016 |
| ctx-lh-paracentral | $\mathrm{GM}_{\mathrm{cort}}$ | 1017 |
| ctx-lh-parsopercularis | $\mathrm{GM}_{\mathrm{cort}}$ | 1018 |
| ctx-lh-parsorbitalis | $\mathrm{GM}_{\mathrm{cort}}$ | 1019 |
| ctx-lh-parstriangularis | $\mathrm{GM}_{\mathrm{cort}}$ | 1020 |
| ⋮ | ⋮ | ⋮ |
| ctx-lh-pericalcarine | $\mathrm{GM}_{\mathrm{cort}}$ | 1021 |
| ctx-lh-postcentral | $\mathrm{GM}_{\mathrm{cort}}$ | 1022 |
| ctx-lh-posteriorcingulate | $\mathrm{GM}_{\mathrm{cort}}$ | 1023 |
| ctx-lh-precentral | $\mathrm{GM}_{\mathrm{cort}}$ | 1024 |
| ctx-lh-precuneus | $\mathrm{GM}_{\mathrm{cort}}$ | 1025 |
| ctx-lh-rostralanteriorcingulate | $\mathrm{GM}_{\mathrm{cort}}$ | 1026 |
| ctx-lh-rostralmiddlefrontal | $\mathrm{GM}_{\mathrm{cort}}$ | 1027 |
| ctx-lh-superiorfrontal | $\mathrm{GM}_{\mathrm{cort}}$ | 1028 |
| ctx-lh-superiorparietal | $\mathrm{GM}_{\mathrm{cort}}$ | 1029 |
| ctx-lh-superiortemporal | $\mathrm{GM}_{\mathrm{cort}}$ | 1030 |
| ctx-lh-supramarginal | $\mathrm{GM}_{\mathrm{cort}}$ | 1031 |
| ctx-lh-frontalpole | $\mathrm{GM}_{\mathrm{cort}}$ | 1032 |
| ctx-lh-temporalpole | $\mathrm{GM}_{\mathrm{cort}}$ | 1033 |
| ctx-lh-transversetemporal | $\mathrm{GM}_{\mathrm{cort}}$ | 1034 |
| ctx-lh-insula | $\mathrm{GM}_{\mathrm{cort}}$ | 1035 |
| ctx-rh-bankssts | $\mathrm{GM}_{\mathrm{cort}}$ | 2001 |
| ctx-rh-caudalanteriorcingulate | $\mathrm{GM}_{\mathrm{cort}}$ | 2002 |
| ctx-rh-caudalmiddlefrontal | $\mathrm{GM}_{\mathrm{cort}}$ | 2003 |
| ctx-rh-cuneus | $\mathrm{GM}_{\mathrm{cort}}$ | 2005 |
| ctx-rh-entorhinal | $\mathrm{GM}_{\mathrm{cort}}$ | 2006 |
| ctx-rh-fusiform | $\mathrm{GM}_{\mathrm{cort}}$ | 2007 |
| ctx-rh-inferiorparietal | $\mathrm{GM}_{\mathrm{cort}}$ | 2008 |
| ctx-rh-inferiortemporal | $\mathrm{GM}_{\mathrm{cort}}$ | 2009 |
| ctx-rh-isthmuscingulate | $\mathrm{GM}_{\mathrm{cort}}$ | 2010 |
| ctx-rh-lateraloccipital | $\mathrm{GM}_{\mathrm{cort}}$ | 2011 |
| ctx-rh-lateralorbitofrontal | $\mathrm{GM}_{\mathrm{cort}}$ | 2012 |
| ctx-rh-lingual | $\mathrm{GM}_{\mathrm{cort}}$ | 2013 |
| ctx-rh-medialorbitofrontal | $\mathrm{GM}_{\mathrm{cort}}$ | 2014 |
| ctx-rh-middletemporal | $\mathrm{GM}_{\mathrm{cort}}$ | 2015 |
| ctx-rh-parahippocampal | $\mathrm{GM}_{\mathrm{cort}}$ | 2016 |
| ctx-rh-paracentral | $\mathrm{GM}_{\mathrm{cort}}$ | 2017 |
| ctx-rh-parsopercularis | $\mathrm{GM}_{\mathrm{cort}}$ | 2018 |
| ctx-rh-parsorbitalis | $\mathrm{GM}_{\mathrm{cort}}$ | 2019 |
| ctx-rh-parstriangularis | $\mathrm{GM}_{\mathrm{cort}}$ | 2020 |
| ctx-rh-pericalcarine | $\mathrm{GM}_{\mathrm{cort}}$ | 2021 |
| ctx-rh-postcentral | $\mathrm{GM}_{\mathrm{cort}}$ | 2022 |
| ctx-rh-posteriorcingulate | $\mathrm{GM}_{\mathrm{cort}}$ | 2023 |
| ctx-rh-precentral | $\mathrm{GM}_{\mathrm{cort}}$ | 2024 |
| ctx-rh-precuneus | $\mathrm{GM}_{\mathrm{cort}}$ | 2025 |
| ctx-rh-rostralanteriorcingulate | $\mathrm{GM}_{\mathrm{cort}}$ | 2026 |
| ctx-rh-rostralmiddlefrontal | $\mathrm{GM}_{\mathrm{cort}}$ | 2027 |
| ctx-rh-superiorfrontal | $\mathrm{GM}_{\mathrm{cort}}$ | 2028 |
| ctx-rh-superiorparietal | $\mathrm{GM}_{\mathrm{cort}}$ | 2029 |
| ctx-rh-superiortemporal | $\mathrm{GM}_{\mathrm{cort}}$ | 2030 |
| ctx-rh-supramarginal | $\mathrm{GM}_{\mathrm{cort}}$ | 2031 |
| ctx-rh-frontalpole | $\mathrm{GM}_{\mathrm{cort}}$ | 2032 |
| ctx-rh-temporalpole | $\mathrm{GM}_{\mathrm{cort}}$ | 2033 |
| ctx-rh-transversetemporal | $\mathrm{GM}_{\mathrm{cort}}$ | 2034 |
| ctx-rh-insula | $\mathrm{GM}_{\mathrm{cort}}$ | 2035 |

(N/A values are due to $T_2$w ISO scans not being acquired on the ENIC 64 mT scanner; see Methods.)

**Table S5**: **Quality control of 64 mT registrations.** Each ultra-low-field scan was rigid-registered to the high-field scan of the corresponding contrast, within participants, using FSL FLIRT (Jenkinson and Smith, 2001). The table contains the number of scans for each registration quality score (1 = perfect / 5 = fail), by ultra-low-field contrast and resolution. We note that readers can assess registration quality by themselves, using publicly available data shared alongside this manuscript, which includes registered scans and segmentations (upon publication).

| | | | score (1 = perfect / 5 = fail) | | | | | |
|---|---|---|---|---|---|---|---|---|
| **scanner** | **contrast** | **resolution** | **1** | **2** | **3** | **4** | **5** | **OK** ($\leq 3$) |
| CNS | $T_1$w | AXI | 11 | 7 | 3 | 2 | 0 | 21 |
| | | COR | 3 | 8 | 6 | 5 | 1 | 17 |
| | | SAG | 11 | 10 | 1 | 0 | 0 | 23 |
| | | ISO | 3 | 10 | 3 | 4 | 3 | 16 |
| | | MRR | 8 | 8 | 5 | 1 | 0 | 22 |
| | $T_2$w | AXI | 7 | 1 | 10 | 3 | 2 | 18 |
| | | COR | 12 | 5 | 4 | 1 | 1 | 21 |
| | | SAG | 4 | 12 | 4 | 2 | 1 | 20 |
| | | ISO | 0 | 1 | 10 | 12 | 0 | 11 |
| | | MRR | 12 | 9 | 1 | 1 | 0 | 22 |
| ENIC | $T_1$w | AXI | 11 | 7 | 3 | 2 | 0 | 21 |
| | | COR | 8 | 9 | 4 | 2 | 0 | 21 |
| | | SAG | 12 | 6 | 5 | 0 | 0 | 23 |
| | | ISO | 4 | 6 | 6 | 4 | 3 | 16 |
| | | MRR | 11 | 6 | 4 | 2 | 0 | 21 |
| | $T_2$w | AXI | 10 | 6 | 4 | 3 | 0 | 20 |
| | | COR | 9 | 12 | 1 | 1 | 0 | 22 |
| | | SAG | 2 | 15 | 4 | 2 | 0 | 21 |
| | | ISO | N/A | N/A | N/A | N/A | N/A | N/A |
| | | MRR | 8 | 15 | 0 | 0 | 0 | 23 |

**Table S6**: **Summary of Dice reliability and correspondence for scans of sufficient registration quality, across contrasts and resolutions.** Global values correspond to the median across 4 global tissue classes. Local values correspond to the median, and first and third quartiles, i.e. Md [$Q_1$, $Q_3$], across 98 SynthSeg+ labels. Values which show a (0.01) difference relative to the main results (with all scans included) are highlighted in bold. (N/A values are due to $T_2$w ISO scans being acquired on one 64 mT scanner only; see Methods.)

| | | **contrast** | **MRR** $T_1$w 1.6×1.6×1.6 $T_2$w 1.5×1.5×1.5 | **ISO** $T_1$w 2.3×2.3×2.3 $T_2$w 2.2×2.2×2.2 | **AXI** $T_1$w 1.6×1.6×5 $T_2$w 1.5×1.5×5 | **COR** $T_1$w 1.6×5×1.6 $T_2$w 1.5×5×1.5 | **SAG** $T_1$w 5×1.6×1.6 $T_2$w 5×1.5×1.5 |
|---|---|---|---|---|---|---|---|
| **Global** | **Dice$_{rel}$** | $T_1$w | 0.85 | 0.81 | 0.84 | **0.83** | 0.83 |
| | | $T_2$w | 0.87 | N/A | 0.84 | 0.85 | 0.86 |
| | **Dice$_{cor}$** | $T_1$w | 0.71 | 0.71 | 0.71 | **0.72** | 0.69 |
| | | $T_2$w | 0.82 | 0.73 | 0.73 | 0.77 | 0.73 |
| **Local** | **Dice$_{rel}$** | $T_1$w | 0.79 [0.77,0.82] | **0.73** [**0.69**,0.77] | 0.75 [0.73,0.79] | 0.74 [**0.70**,0.77] | 0.75 [**0.72**,0.78] |
| | | $T_2$w | 0.81 [**0.78**,0.84] | N/A | 0.76 [0.71,0.79] | 0.77 [0.74,0.81] | 0.77 [0.75,0.81] |
| | **Dice$_{cor}$** | $T_1$w | 0.61 [0.53,0.70] | 0.59 [0.52,0.69] | 0.61 [0.54,0.66] | **0.60** [**0.52**,0.69] | 0.56 [**0.49**,**0.65**] |
| | | $T_2$w | 0.74 [0.68,0.79] | 0.65 [0.57,0.72] | **0.61** [**0.51**,**0.70**] | 0.67 [**0.61**,0.73] | 0.59 [0.54,0.66] |
